# Loss-of-Function genetic Screen Unveils Synergistic Efficacy of PARG Inhibition with Combined 5-Fluorouracil and Irinotecan Treatment in Colorectal Cancer

**DOI:** 10.1101/2025.03.29.644889

**Authors:** Cristina Queralt, Cristina Moreta-Moraleda, Marta Costa, Ferran Grau-Leal, Jeannine Diesch, Carla Vendrell-Ayats, Eva Musulén, Cristina Bugés, José Luis Manzano, Sara Cabrero, Johannes Zuber, Marcus Buschbeck, Sonia Forcales, Eva Martínez-Balibrea

**Author notes:** Correspondence to: Dr. Eva Martínez-Balibrea, Germans Trias i Pujol Research Institute (IGTP) and Catalan Institute of Oncology, Ctra. De Can Ruti, camí de les escoles s/n, 08916 Badalona, Spain. and Dr. Sonia Forcales, University of Barcelona, c/Feixa llarga s/n, L’Hospitalet de Llobregat, 08907, Spain. JD: EM: CB, JLM: JZ: MB: SF, CM.

## Abstract

Colorectal cancer (CRC) remains a major global health concern, partly due to resistance to therapy and the lack of new effective treatments for advanced disease. The combination of 5-Fluorouracil (5FU, a thymidylate synthase inhibitor) and irinotecan (a topoisomerase 1 inhibitor) is widely used in first-line and subsequent treatments. This study aimed to identify novel therapeutic targets to enhance combinatorial therapy, improving treatment efficacy and durability of response. We performed a loss-of-function screen using HT29 CRC cell line and a retroviral library containing 7296 shRNAs targeting 912 chromatin genes. Cells were then treated with 5FU and SN38 (the active metabolite of irinotecan) or left untreated for 4 weeks. Genes enriched in resistant clones were identified through next-generation sequencing. Among candidate genes, PARG was selected for functional validation. CRISPR/Cas9-mediated knockout (HT29 PARG-KO) resulted in increased global poly(ADP-ribosyl)ation after 5FU and SN38 treatment. PARG depletion led to reduced cell viability and increased apoptosis, particularly after 5FU exposure. Pharmacological PARG inhibition (PDD00017273) synergized with 5FU and SN38 across three CRC models (HT29, DLD1, HT115). *In vivo*, HT29 PARG-KO xenografts were more sensitive to 5FU. Immunohistochemical analysis of 170 CRC patient tumors revealed that positive PARG expression correlated with poor response to 5FU + Irinotecan, increased liver metastases, and worse long-term survival. Our findings highlight PARG as a promising therapeutic target for CRC, where its inhibition enhances the efficacy of standard chemotherapy.

## 1. Background

Colorectal cancer (CRC) is the most common tumour in both sexes worldwide, with serious social and economic repercussions. In recent years, the increasing incidence in individuals under 50 years of age has placed it under the spotlight. Overall survival (OS) at 5 years is around 65%, although in metastatic disease this does not exceed 15%. The treatment of choice in these cases has remained largely unchanged for more than two decades, consisting of combinations of classic chemotherapy (fluoropyrimidines, oxaliplatin, irinotecan) with an antiangiogenic agent or an EGFR inhibitor (only in patients whose tumours do not carry mutations in the RAS oncogenes). This approach has resulted in objective responses in approximately 50% of cases, with acquired resistance being the main cause of tumor progression in responders. In light of this, it is imperative to identify new potential therapeutic targets that offer treatment alternatives, either in monotherapy or in combination with existing drugs.

The approval of the combination therapy comprising 5-fluorouracil (5FU), leucovorin (LV), and irinotecan (FOLFIRI) for metastatic CRC in 2000 marked a significant advancement in treatment efficacy over the standard 5FU and LV regimen (1). Since its approval, FOLFIRI (and other schedules combining 5FU and irinotecan) has emerged as one of the cornerstones of metastatic CRC therapy, alongside the alternative regimen of 5FU, LV, and oxaliplatin. 5FU represents an uracil analogue wherein a hydrogen atom in the carbon 5 position of the pyrimidine ring is substituted with a fluorine atom. Following a cascade of enzymatic transformations, 5FU undergoes conversion into active metabolites including fluorodeoxyuridine monophosphate (FdUMP), fluorodeoxyuridine triphosphate (FdUTP), or fluorouridine triphosphate (FUTP). These metabolites exert inhibitory effects on thymidylate synthase (TS) upon integration into DNA or RNA. Such inhibition disrupts the synthesis of dTMP, leading to perturbations in the deoxynucleotide pool, particularly altering the ratio of dATP to dTTP. Consequently, this dysregulation profoundly impairs DNA synthesis and repair mechanisms, culminating in cellular demise. Irinotecan, or CPT-11, is a water-soluble derivative of camptothecin, a natural plant alkaloid. Upon metabolism, irinotecan gives rise to SN38, which serves as a potent inhibitor of topoisomerase I (*TOP1*). SN38 specifically binds to the DNA-topoisomerase 1 complex, obstructing DNA replication and inducing double-stranded DNA breaks, thereby triggering apoptotic cell death. These actions predominantly occur during the DNA synthesis phase (S phase), rendering irinotecan a cycle-specific cytotoxic agent.

Chromatin regulators are proteins and protein complexes that govern the structure and accessibility of chromatin. Their principal function lies in orchestrating gene expression modulation by finely tuning DNA accessibility to transcription factors and other regulatory proteins. This pivotal role extends to DNA repair processes, where they wield influence over the recruitment and activity of repair machinery at sites of DNA damage. Therefore, chromatin regulators can play a significant role in modulating the efficacy of 5FU plus irinotecan treatment in CRC with some of them being proposed as biomarkers of response and as possible therapeutic targets whose inhibition could result in synergistic treatments (2). Using a loss-of-function (LOF) genetic screen specifically designed to silence genes coding for chromatin regulatory factors, we aimed to identify potential therapeutic targets whose combination with 5FU + irinotecan results in synergistic treatment in CRC.

## 2. Materials and methods

### 2.1. Cell culture

HT29, DLD1 and HT115 human CRC cell lines were obtained from the American Type Culture Collection. Supplementary Table ST1 A shows the list of mutations for each of the cell lines according to the data published in the <u>Cellosaurus</u> and <u>COSMIC</u> databases. Platinum-E (PLAT-E) and HEK293T cells were used for retro- and lentiviral production, respectively. Cells were grown at 37 °C and 5% CO_2_ in McCoy’s 5A medium GlutaMAX (Lab Clinics) (HT29), RPMI 1640 medium (LabClinics) (DLD1 and HT115) and DMEM/F12 medium (Invitrogen) (Platinum-E (PLAT-E) and HEK293T) supplemented with 10% fetal bovine serum (FBS) (Reactiva) and 1% Penicillin-Streptomycin-Amphotericin B (Invitrogen). Additionally, RPMI and DMEM/F12 media were supplemented with 10 mM glutamine (Invitrogen), and 10 mM HEPES (Thermo). Cells were periodically tested for Mycoplasma contamination and authenticated by short tandem repeat profiling using in-house methods.

### 2.2 Drugs

5FU and irinotecan were obtained from leftover vials prepared for patients at the pharmacy of the Catalan Institute of Oncology. SN38, the active metabolite of irinotecan, and the PARG inhibitor PDD00017273 (hereafter referred to as PDD) were purchased from MERCK and TOCRIS Bio-Techne, respectively. Both drugs were reconstituted in DMSO to obtain a stock concentration of 5 mM and 10 mM, respectively, and stored at −20 °C. Further dilutions of each drug were made in culture medium.

### 2.3. Lentiviral production

This procedure was employed to introduce the ECO receptor (pWPXLd-rtTA3-IRES-EcoRec-PGK-Puro construct) for biosafety reasons and to generate a knockout (KO) model for PARG using the CRISPR/Cas9 technique, both in the HT29 cell line. The viral particles were generated by adding to HEK293T cells the packaging plasmids psPax2 and pCMV-VSV-G (both from Addgene) following Moore vector ratio (1:1:1) using Lipofectamine 2000 (Invitrogen) according to manufacturer’s protocol. After 48h of transfection, viral supernatant was filtered in 0.45 µm filters (Merck-Millipore) and added to HT29 cells at 70% of confluence in the presence of 8 µg/µL of Polybrene (Merck-Sigma Aldrich). 72 hours later, HT29 cells were selected for at least one week with 0.35 µg/mL puromycin (Merck-Sigma Aldrich). The introduction of the Eco receptor into our cells was confirmed through RT-qPCR (Supplementary Figure S1). The primers used are listed in Supplementary Table ST2.

### 2.4. CRISPR/Cas9 PARG-KO

HT29 cell line was infected with lentiviral particles containing a pool of three different gRNA for the PARG gene (pLentiCRISPR v2; GenScript) generated independently following the previously described protocol (Supplementary table ST3). After 48 hour-production of individual gRNA lentivirus, independently viral supernatants were filtered and mixed in a 1:1:1 proportion doing a pool. 1/40 dilution of the viral mix was added to HT29 cells at 70% of confluence in a proportion of 1:3 of total plate medium volume. At 4- and 24-hours post-infection 1:3 of fresh media was added. After one week of puromycin selection, single-cell sorting was performed by FACSAria II (BD Biosciences) to isolate homogenous edited cells within the mixed pooled edited population. The clonal edition was confirmed by Sanger sequencing (See primers in Supplementary Table ST2). A negative Non-Target control (NTC) (NonTargetingControl Guide for Human, GenScript) was also infected in the HT29 cell line following the same procedure. The resulting cell lines were named HT29 PARG-KO and HT29 NTC, respectively.

### 2.5. Retroviral production

This method was used to infect HT29 EcoR cells with the shRNA hEPI9 library and to introduce to HT29 cells the individual shRNA for validation. The retroviral particles were generated by adding to Plat-E cells the shRNA hEPI9 library or individual shRNA (both cloned in pMSCV-LENC vector following standard cloning techniques) using Lipofectamine 2000 (Invitrogen), following manufacturer’s protocol. After 72 hours, viral supernatant was collected and filtered using 0.45 µm filters (Merck-Millipore). A proportion of 1:3 of total plate medium volume of viral supernatant was added to cells at 70% of confluence in the presence of 8 µg/µL of Polybrene (Merck-Sigma Aldrich). After 4 hours post-infection, up to 2:3 of fresh medium was added. 72 hours post-infection, cells were selected with 600 µg/mL of Geneticin (Thermo) for at least 1 week. The percentage of cell infection was measured by Flow Cytometry using mCherry cells positivity in a LSRFortessa SORP Flow Cytometer (BD Biosciences). For the subsequent individual validation of PARG, selected shRNAs and a non-human shRNA control (shRenilla coming from firefly) was also infected following the same procedure.

### 2.6. Retroviral shRNA hEPI9 library infection and LOF Screening

The miR-E-based short hairpin library, hEPI9, comprises 7296 shRNAs targeting 912 chromatin genes (with 8 shRNAs per target) including both positive and negative shRNA controls. The functions of hEPI9 targeted genes were represented in Supplementary Figure S2. This library was kindly provided by Dr. Johannes Zuber (2,3). A retroviral infection efficiency of 1% was observed in previous experiments using HT29 EcoR cells. Since the hEPI9 library contains close to 7,300 shRNAs and we wanted to achieve a 1,000X representation of each one, 730 x 10^6^ HT29 EcoR cells were infected with the library retroviral supernatant following the protocol described above. After selection, cells were treated or not using the IC_20_ of FUIRI (0.1 µM of 5FU + 0.055 nM of SN38), 24 hours after seeding. The next day, the medium was changed, and cells were allowed to grow for 72 hours. This schedule was repeated four consecutive times for three weeks. After 21 days of treatment, cells were collected and prepared for sequencing.

### 2.7. Drug-response analysis by the 3-(4,5-dimethylthiazol-2-yl)-2,5-diphenyltetrazolium bromide (MTT) assay

To calculate the inhibitory concentrations (IC) of different drugs and combinations, we used the MTT assay (Roche) as previously described (3). Cells were seeded in 96-well microtiter plates (Nunc) at densities of 1,500 cells per well for HT29 and DLD1, and 5,000 cells per well of HT115 cell line, incubated for 24 hours and treated with a range of serial drug concentrations for 24 hours. Subsequently, the drug-containing media was removed and replaced with fresh media, allowing the cells to grow for 72 hours. MTT was then added to each well and, following incubation, the resulting formazan crystals were solubilized using a mixture of 0.02 M HCl and 20% SDS (50% v/v). The absorbance was measured at 570 nm using the SPECTROstar® Nano spectrophotometer (BMG Labtech). The drug concentrations corresponding to each fraction of survival (ranging from 10% to 90% cell viability) were determined using the median-effect line method. The maximum concentration of DMSO (vehicle) used was 0.5%, which did not affect cell proliferation in our experiments.

For the LOF screen, we established the FUIRI IC_20_: HT29 EcoR cells were treated with serial dilutions ranging from 1/6 to 1/100 of 5FU and SN38 IC_50_ in 1:1 proportion for 24 hour-treatment following the MTT assay. FUIRI IC_20_ (0.1 µM of 5FU + 0.055 nM of SN38) was determined after four consecutive 24-hour treatments (Supplementary Figure S3) mimicking the schedule followed for LOF screen.

### 2.8. Analysis of combined drug effects

Cells were seeded at the corresponding cell density in a 96-well plate, incubated for 24 hours and treated with a range of serial dilutions of the previously established IC_50_ values for 5FU and SN38 for 72 hours. FUIRI combinations (5FU + SN38) were prepared by mixing 5FU and SN38 at a 1:1 ratio of their IC_50_ values, and serial dilutions were made from 1/16 to 4 times the IC_50_. Additionally, PDD was added at final concentrations of either 1 µM or 5 µM. To evaluate the combined drug effects, a combinational index (CI) was calculated using the CompuSyn software v.1, following the Chou and Talalay method. The CI was interpreted as follows: CI = 1 indicates an additive effect, CI < 1 indicates synergism, and CI > 1 indicates antagonism.

### 2.9. Next Generation Sequencing Solexa Technology

DNA was extracted using the Phenol-Chloroform-Isoamylalcohol protocol and sent for sequencing by NGS (Illumina HiSeq2500 platform) to Dr. Zuber’s lab. The sequencing results were analyzed using the R package EdgeR (R Core Team 2019) as described (3), which calculated log-fold change (logFC) values, counts per million, significance values (p-values), and false discovery rates (FDR). Genes were selected if at least 6 out of 8 hairpins exhibited consistent behavior in the same direction and were ranked on a gene-by-gene basis according to mean logFC, p-value, and FDR values using the Roast function.

### 2.10. Gene expression by Real Time-quantitative PCR (RT-qPCR)

RNA was extracted from cultured cells using the Maxwell® 16 LEV simplyRNA Cells Kit (Promega) according to the manufacturer’s protocol. cDNA synthesis was performed with 1 µg of RNA using the SuperScript™ IV First-Strand Synthesis System (Thermo) following the manufacturer’s instructions. Quantitative PCR (qPCR) was carried out using pre-designed IDT™ assays and SYBR® Green master mix (Roche) in a LightCycler® 480 II instrument (Roche) (Supplementary Table ST2). To ensure the absence of genomic DNA contamination, RT minus controls (no reverse transcriptase) were included for each sample, with contamination acceptable if below 5% of the total signal. All qPCR reactions were conducted in triplicates to ensure reproducibility and reliability of the results. Relative gene expression levels were calculated using the 2^(-ΔΔCt) method, with gene expression normalized to the housekeeping gene β-actin (IDT™) and cells transfected with the control vector used as reference samples to determine relative expression levels. Data were analyzed using LightCycler® 480 software version 1.5 (Roche) for threshold cycle (Ct) determination.

### 2.11. Propidium iodide-based cell proliferation assay

Propidium iodide (PI) was used as a dye that penetrates only damaged cellular membranes. NTC and PARG-KO cells were seeded in 24-well plates and after 24 hours, PI (Merck-Sigma Aldrich) was added to a final concentration of 90 µM. Protected from light, the non-viable cell population was detected by the first fluorescence measurement (excitation and emission wavelengths were 560 nm and 654 nm, respectively). Then, triton x-100 (Merck-Sigma Aldrich) was added at a final concentration of 1.7%. After 5 minutes of incubation at room temperature, the second measurement was obtained at the same wavelengths to determine the total cell population count. The number of viable cells was calculated by subtracting the first measurement from the second, at each tested timepoint. The fluorescence was measured with the Thermo Scientific Varioskan Flash (Thermo) plate reader.

### 2.12. Cell viability assay

Cells were seeded in 6-well plates to confirm the effect of gene knockdown (KD) or KO upon treatment. Cells were treated for 24 hours with serial dilutions (1/100, 1/50, 1/6, and 2X) of the combination of FUIRI IC_50_ as previously established by the MTT assay. After 24 hours of treatment, the drug-containing medium was removed, and the cells were allowed to grow for an additional 72 hours. Subsequently, cells were collected along with the corresponding supernatant and centrifuged. Pellets were resuspended in 1X PBS and stained for 30 minutes at 37°C with 10 µM DiOC (Thermo). Following two washes with 1X PBS, cells were stained for 15 minutes with 3 µM DAPI (Merck-Sigma Aldrich) in 1X PBS. Fluorescence levels were measured by flow cytometry using the LSR Fortessa SORP Flow Cytometer (BD Biosciences) and analyzed with BD FACSDivaTM software (BD Biosciences).

### 2.13. Sanger DNA sequencing

DNA was extracted using the Maxwell® RSC Cultured Cells DNA Kit (Promega) according to the manufacturer’s protocol. 50ng of DNA were then amplified by PCR with Phusion™ High-Fidelity DNA Polymerase (2 U/µL) (Thermo) and specific primers (IDT™) for 30 cycles (Supplementary Table ST2). Following amplification, the PCR product was verified for correct band size by 1% agarose gel electrophoresis. The product was purified using ExoSAP-IT™ PCR Product Cleanup Reagent (Thermo) as per the manufacturer’s instructions. Finally, the samples were sent for Sanger sequencing at Macrogen Inc.

### 2.14. Apoptosis assays

HT29 NTC and PARG-KO cells were treated 24 hours after seeding in 6-well plates with serial dilutions (1/18, 1/3, 1X, and 2X) of the combination of individual IC_50_ values for 5FU and SN38 or the FUIRI combination. After 72 hours of treatment, cells were collected along with the supernatant and centrifuged. Following two washes with cold 1X PBS, the cell pellets were resuspended in 1X Binding Buffer (BD Pharmingen™) and stained for 15 minutes at room temperature, protected from light, with 4 µl of FITC Annexin V and 3 µl of Propidium Iodide (BD Pharmingen™). Prior to measuring fluorescence levels using flow cytometry with the LSR Fortessa SORP Flow Cytometer (BD Biosciences), 400 µl of 1X Binding Buffer (BD Pharmingen™) was added to the cells. The results were analyzed with BD FACSDiva™ software (BD Bioscience).

### 2.15. Total protein and acidic histone extraction

Total protein extraction began with the homogenization of dry frozen cell pellets in RIPA buffer (PBS; NP-40 1%; Na deoxycholate 0.5%; SDS 0.1%; EDTA 1 mM; NaF 50 mM; NaVO3 5 mM) supplemented with a cocktail of EDTA-free protease inhibitors (Roche) using the gentleMACS Dissociator system (Miltenyi Biotec). After a 15-minute incubation at 4°C, the samples were centrifuged at maximum speed for 15 minutes at 4°C. The aqueous supernatant containing the total protein extract was quantified using the Pierce™ BCA Protein Assay Kit (Thermo) according to the manufacturer’s protocol. For histone isolation, the remaining pellets were resuspended in 0.2 M HCl, homogenized with the gentleMACS Dissociator system (Miltenyi Biotec), rotated at 4°C for 15 minutes, and then neutralized by adding Tris-HCl (1 M; pH 8).

### 2.16. Western Blot (WB)

Fifty micrograms of total protein or 25 µl of histone proteins were loaded onto NuPAGE™ 10% Bis-Tris Midi Protein Gels (Invitrogen) and run using MES SDS running buffer 1X (Invitrogen), at 180 V for 1 hour. Gels were dry transferred to PVDF membranes using the iBlot 2 Dry Blotting System (Thermo). After transferring, membranes were blocked for at least 1 hour at room temperature with Intercept® (TBS) Blocking Buffer (LI-COR, Invitrogen). Membranes were then incubated with specific primary antibodies overnight at 4°C. The following day, after several washes with 1X TBS-Tween 0.1%, membranes were incubated with secondary LI-COR antibodies for 1 hour at room temperature, protected from light. After several washes with 1X TBS-Tween 0.1%, membranes were scanned and analyzed using the LI-COR Odyssey 9120 Digital Imaging System. Band quantification was performed using Image Studio™ Software and normalized by comparing phosphorylated proteins to their non-phosphorylated counterparts or to a housekeeping protein such as α-tubulin. The antibodies used for WB were: 1/1,000 PARG mouse antibody (OTI6F4) (NBP2-46320; Novus); 1/20,000 α-tubulin mouse antibody (T6074; Sigma); 1/1,000 Anti-Histone H2A.X rabbit antibody (AB11175; Abcam); 1/1,000 Anti-phospho-Histone H2A.X (Ser139) mouse antibody clone JBW301 (05-636; Merck); and 1/1,000 PAR mouse antibody (AM80; Sigma).

### 2.17. Immunofluorescence

HT29 NTC and HT29 PARG-KO cells were seeded in LabClinics chambers. After 24 hours, cells were treated with a 1:3 dilution FUIRI IC_50_, adding PDD at 5 µM alone or in combination with 5FU and FUIRI for 24 hours. Following treatment, the media was replaced with formaldehyde (4% in TBS) for fixation. Permeabilization was performed using 0.2% Triton X-in 1X TBS, and blocking was done with 1% BSA in 1X TBS for 1 hour. Cells were then incubated with a 1:100 dilution of γH2A.X antibody (05-636; Merck) in freshly prepared 1% BSA-1X TBS for 90 minutes. After several washes, cells were stained with a 1:1,000 dilution of fluorophore-conjugated Goat anti-Mouse Alexa 555 secondary antibody (A-21422; Invitrogen) in freshly prepared 1% BSA-TBS for 30 minutes, protected from light. Following additional washes, cells were stained with a 1:20,000 dilution of 0.5 mg/ml DAPI (Merck-Sigma Aldrich) for 5 minutes to visualize the nucleolus. Slides were mounted with ProLong™ Gold Antifade (Thermo). Images were captured using the Micro Axio Imager M2 Apotome (Zeiss) and the total γH2AX foci/field were quantitated by the Find Maxima feature of ImageJ software. At least 200 nuclei were analyzed per sample.

### 2.18. *In vivo* experiments

Five-week-old athymic nude Foxn1^nu^ male and female mice were purchased from Envigo and acclimated in a temperature- and humidity-controlled room at the Center for Comparative Medicine and Bioimage (CMCiB) on a 12-hour light-dark cycle for one week prior to the experiment. Mice were housed under specific pathogen-free conditions and monitored according to strict welfare protocols adhering to the 3Rs principle. A preliminary tumor growth and toxicity test was conducted before to evaluate treatment efficacy. Tumor growth was assessed by subcutaneously injecting 3 × 10C of HT29 NTC or PARG-KO cells into the left flank of each mouse (100 µl) in McCoy’s 5A medium GlutaMAX (LabClinics) mixed with Matrigel Matrix Basement Membrane (Corning) at a 1:1 ratio into 10 mice per group (5 females and 5 males each). Tumor growth was measured twice a week using a digital caliper. Tumor volume (TV) was calculated using the formula V = 1/2 × length (mm) × [width (mm)]² over 21 days. Treatment toxicity was assessed as follows: ten mice were treated intraperitoneally (IP) once a week for four weeks with either PBS 1X (n = 2), 50 mg/kg CPT-11 (n = 2), 50 mg/kg 5FU (n = 2), or a combination of 50 mg/kg 5FU + 50 mg/kg CPT-11 (FUIRI) (n = 4). Toxicity was assessed by weekly monitoring of body weight. A suspension of 2.5 ·10^6^ HT29 NTC or HT29 PARG-KO cells in McCoy’s 5A medium GlutaMAX (LabClinics) mixed with Matrigel Matrix Basement Membrane (Corning) at a 1:1 ratio was injected subcutaneously into the left flank of each mouse (100 µl). Tumor growth was measured twice a week using a digital caliper, and TV was calculated with the formula previously described. Once tumors reached 100 mm³, mice were treated intraperitoneal (IP) with vehicle (PBS 1X), 50 mg/kg 5FU, 50 mg/kg CPT-11, or FUIRI (50 mg/kg 5FU + 50 mg/kg CPT-11) once per week for four weeks, according to their treatment group. Relative Tumor Volume (RTV) was determined by the ratio of each measurement’s TV to the TV on the first day of treatment for each group. Tumor Growth Inhibition (TGI) was calculated using the formula TGI (%) = [1 − (RTV of the treated group) / (RTV of the control group)] × 100 (%). At the end of the experiment, mice were fully anesthetized and euthanized by cervical dislocation. The Animal Experimentation Commission of the Catalan Government approved all *in vivo* procedures under project reference 11730.

### 2.19. PARG Immunohistochemical analysis

We used 6 Tissue Microarrays (TMAs) constructed from the FFPE primary tumors of 170 metastatic CRC patients who received either irinotecan (180 mg/m^2^ on day 1) + 5FU (400 mg/m^2^ bolus and 600 mg/m^2^ 22-h infusion) + leucovorin (LV) (200 mg/m^2^ on days 1-2) (FOLFIRI) every 2 weeks or irinotecan (80 mg/m^2^) + 5FU (250 mg/m^2^ 48-h infusion) (FUIRI) weekly, as first-line treatment, in the framework of a clinical study (4). All patients signed informed consents to participate in the study. Patient characteristics are in Table 1.

**Table 1.** PARG inhibitors

| **Name** | **IC50** | **Characteristics** | **Ref** | **Model** |
| --- | --- | --- | --- | --- |
| **Gallotannin** | 16,8 μM | Low activity<br>Off-target effects | Tsai et al., 1991 | Preclinic |
| **Rhodamine-based PARG inhibitors** | 1-6 μm | Selective<br>Not cell-permeable | Finch et al., 2012 | Preclinic |
| **GPI** | 1,7 μM | Low activity<br>Off-target effects | Lu et al., 2003 | Preclinic |
| **ADP-HPD** | 120 nM | Selective<br>Lack of cell permeability | Slama et al., 1995 | Preclinic |
| **PDD00017273** | 26 nM | Selective<br>Cell permeable<br>Low bioavailability | James et al., 2016 | Preclinic |
| **COH34** | 0,37 nM | Selective<br>High efficiency of inhibition<br>Cell permeable | Chen and Yu, 2019 | Preclinic |
| **JA2131** | 400 nM | Selective<br>Cell permeable<br>Could be modified to increase inhibition potency | Houl et al., 2019 | Preclinic |
| **IDE161** | N/A | Selective<br>High efficiency of inhibition<br>High preclinical tolerability profile | Monah et al., 2023 | Phase 1<br>NCT05787587 |
| **ETX-19477** | Low nM | Selective<br>High efficiency of inhibition<br>Well tolerated<br>Robust dose-dependent in vivo anti-tumour activity | Holleran et al., 2024 | Phase 1<br>NCT06395519 |

Most commonly used PARG inhibitors. All of these inhibitors have been evaluated in vitro to assess their efficacy. Notably, the last two inhibitors are currently being tested in Phase I clinical trials for solid tumours including CRC.
N/A: not applicable.

Five µm thick TMA sections were analyzed using standard IHC techniques. PARG expression was detected using a PARG mouse antibody (OTI6F4) (NBP2-46320; Novus). The process was performed automatically using a Ventana BenchMark ULTRA machine (Roche). An external positive control was included on each slide. Immunostaining was independently evaluated by two pathologists. PARG expression was assessed using a 4-level scoring system: null (0), mild (1+), medium (2+), and strong (3+).

### 2.20. Statistical analysis

Statistical analyses of the experimental data were performed using the GraphPad Prism software (version 8). The specific statistical test used for each experiment is specified in the corresponding figure legend. Data visualization includes results from at least three independent experiments, with figures showing mean values along with either standard deviations (SD) or standard error of the mean (SEM) as indicated. To evaluate significant differences between groups, Student’s t-Test analysis and twoCway analysis of variance (ANOVA) were used, when apropriate. Statistical significance was defined as p < 0.05. For *in vivo* experiments Linear Mixed Effects (LME) model was used to analyze the effect of treatments. Data from the patients study were analyzed using PASW Statistics (version 18 for Mac), with the tests used specified in each figure caption. Overall survival data are represented as Kaplan-Meier curves and the p value was calculated using the Log-Rank test.

## 3. Results

### 3.1. Genetic LOF Screening Identifies PARG as a Potential Combinatorial Drug Target in CRC

We used the shRNA hEPI9 library to perform a genetic LOF screening in a pooled approach to identify chromatin-related genes that synergize with FUIRI treatment. An optimized mirE backbone was used to clone all shRNAs, increasing KD efficiency of target genes after single-copy integration (5,6) (Figure 1A). As a cellular model, we chose the human CRC cell line HT29, which is well-characterized in our laboratory (7–9). According to the COSMIC and Cellosaurus databases, these cells harbour mutations in key CRC-associated genes *APC* (p.E853*), *BRAF* (p.Val600Glu), *PIK3CA* (p.Pro449Thr), and *TP53* (p.Arg273His), and are classified as CMS3 (10,11) (Supplementary Table ST1 A and B). The selected shRNAs target human chromatin-related genes, so the cells were modified by introducing a murine ecotropic receptor via lentiviral infection (Supplementary Figure S1). This strategy prevents harmful effects for the researcher, as these cells can only be infected by murine retrovirus that recognizes this receptor. For the LOF screening, HT29-EcoR (hereafter HT29) cells were infected with the hEPI9 shRNA library at low viral titter to favor single-copy integration. After geneticin selection, cells were treated four consecutive times over three weeks with a combination of 5FU and SN38 at their corresponding IC_20_ (0.1 µM + 0.055 nM, respectively) (Supplementary Figure S3). In mild treatments like IC_20_, shRNAs that drop out can pinpoint the most vulnerable genes. After this period, the pool of shRNAs integrated into the genomic DNA was compared between treated and untreated cells via NGS (Figure 1B). Potential candidate genes were selected based on the combination of different parameters such as the trend consistency of the shRNA (genes that showed the down representation of at least five out of eight shRNAs compared to untreated cells), the mean of the log fold change of the eight shRNA (the more negative values indicated higher sensitivity to treatment), the mixed p value of the eight shRNA (p < 0.1) and the FDR (< 0.5).

**Figure 1.**
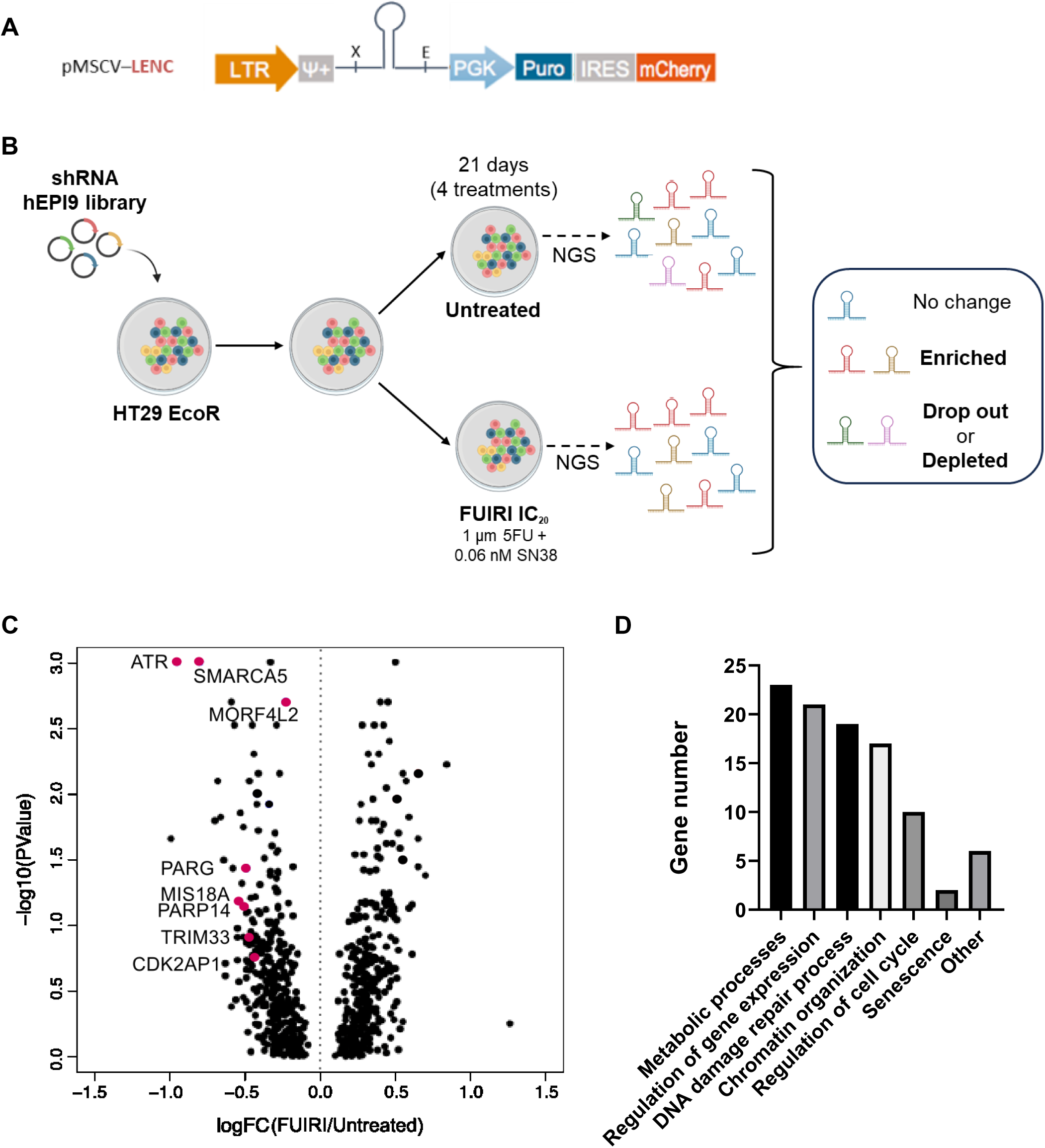
LOF screening design and results. A) pMSCV vector. Diagram illustrating the elements of the pMSCV vector including a long terminal repeat (LTR) for integration, an extended retroviral packaging signal (Ψ), a shRNA integration site, a constitutive phosphoglycerate kinase promoter (PGK), a puromycin resistance gene (Puro) and an internal ribosomal entry site (IRES) for transcription and red fluorescent protein (mCherry). B) LOF screening timeline. The HT29 EcoR cell line was infected with the hEPI9 library and treated with the IC_20_ of FUIRI for four consecutive treatments over 21 days. NGS was used to compare the shRNA representation between treated and untreated cells. C) Volcano plot of candidate hits from the LOF screening. Of the 912 chromatin factors tested, 352 genes were significantly depleted, potentially indicating an association with FIUIRI sensitivity (left side of the volcano plot). In contrast, 131 genes were enriched suggesting a possible association with a lack of response to FUIRI treatment (right side of the volcano plot) based on logFC. Genes associated with FUIRI sensitivity, validated individually, are represented by red dots. D) Gene Ontology (GO) analysis of the biological processes related to the depleted genes. GO analysis was performed using the PANTHER Overrepresentation Test, applying Fisher’s Exact Test without any statistical correction.

Using this approach, we identified 352 genes whose shRNAs were significantly depleted and 131 genes whose shRNAs were enriched, represented in the volcano plot in Figure 1C. The 24 top drop-out genes identified are listed in Supplementary Table ST4. A decrease in shRNAs in treated cells indicated that KD of the corresponding genes was not tolerated, suggesting that the gene products are essential for survival in the presence of FUIRI treatment. A Gene Ontology (GO) analysis was performed to identify the most common functions of the potential candidate genes (Figure 1D). Genes associated with sensitivity to FUIRI treatment were primarily involved in the regulation of metabolic processes and gene expression, regulation of DNA damage response and chromatin remodelling as well. We selected eight genes for individual validation, by generating a polyclonal HT29 cell line with a stable single KD for each gene. The viability of these KD cells was assessed after a 24-hour FUIRI treatment across a range of concentrations, starting from low doses similar to the used in the LOF screening and increasing up to, and beyond, the previously calculated IC_50_, to rule out potential targets. The efficiency of the KD was assessed by measuring the decrease in RNA and protein levels in HT29 EcoR cells infected with each specific shRNA. The best results were obtained *for ATR, MORF4L2, SMARCA5* and *PARG* (Supplementary Figure S4A and S4B). Poly(ADP-Ribose) Glycohydrolase (PARG) is the primary enzyme responsible for degrading poly(ADP-ribose) (PAR). Following DNA damage, PARG removes PAR chains from proteins involved in DNA repair, facilitating this process. Moreover, since PAR degradation generates AMP, PARG plays a role in cellular energy balance and other key processes, including cell metabolism, apoptosis, genomic stability and transcription regulation. While pharmacological inhibitors are already available for both ATR (12–14) and PARG (15–18), due to the limited scientific evidence on the potential synergism between PARG inhibitors and the FUIRI combination—unlike ATR inhibitors—we decided to continue our study with PARG.

In summary, our genetic LOF screen of chromatin regulators identified several genes affecting sensitivity to FUIRI in HT29 cells, specifically highlighting PARG as a potential drug target for combinatorial treatments.

### 3.2. *PARG* Editing Enhances DNA Damage, Sensitivity, and Apoptosis Induction Following FUIRI and, More Pronouncedly, 5FU Treatment

To perform further functional studies, we established an alternative PARG-KO cell model utilizing CRISPR/Cas9 technology to abolish *PARG* expression. While both shRNA and CRISPR/Cas9 systems are valuable for functional investigations, the enhanced stability and reduced likelihood of off-target effects of the latter influenced our decision to use this model for subsequent experiments. The *PARG* gene was successfully edited in the exon 7 region (Figure 2A), resulting in undetectable protein staining by WB (Figure 2B). Using the same concentrations of the FUIRI combination, we observed a comparable or even greater reduction in post-treatment cell viability (Figure 2C) compared to the KD model. However, the abolition of *PARG* expression had no impact on cell proliferation in the absence of treatment (Supplementary Figure 5).

**Figure 2.**
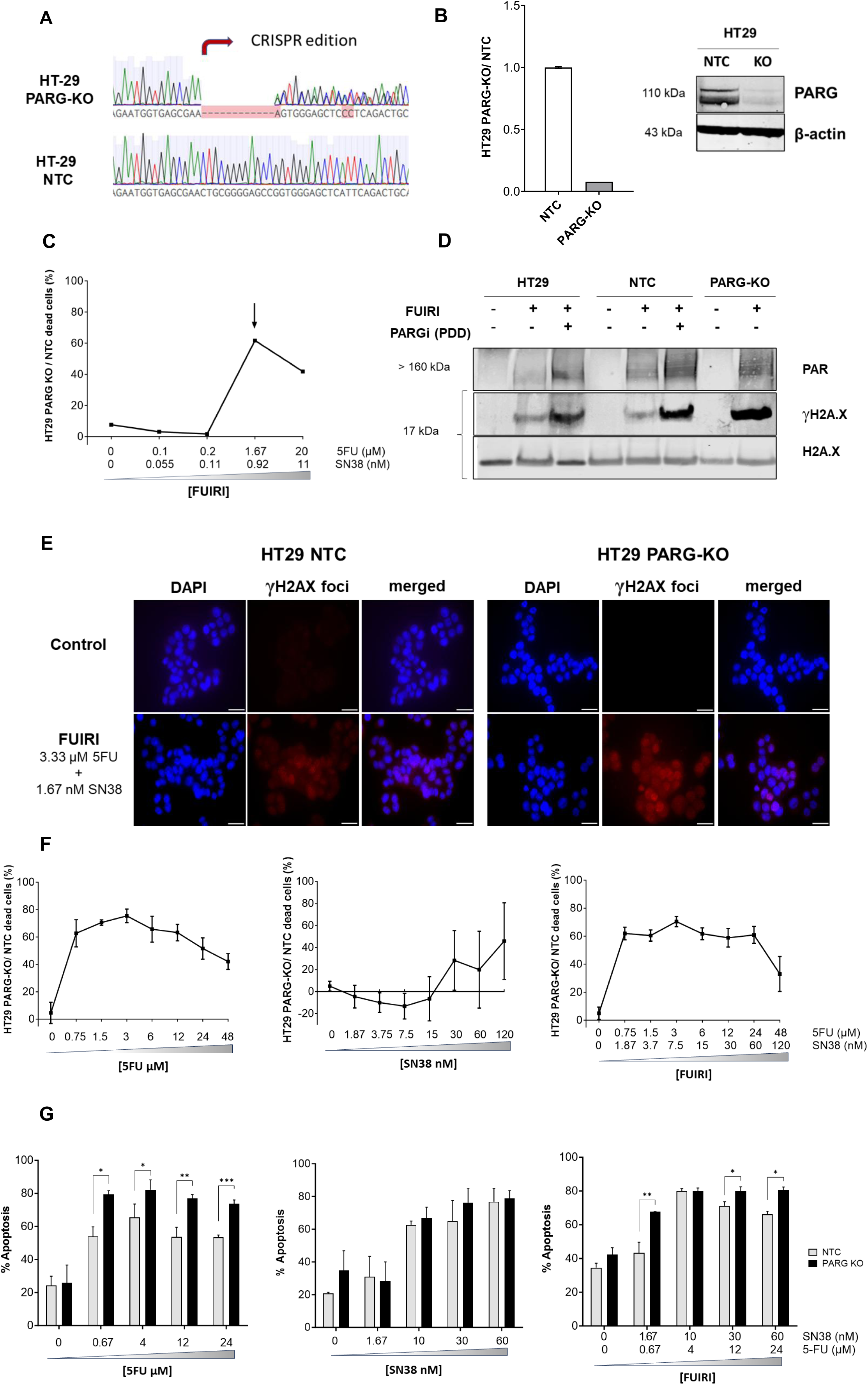
CRISPR/Cas9-based PARG gene editing and functional validation in HT29 cells. A) Representative Sanger sequencing results of DNA from CRISPR/Cas9-edited PARG HT29 clones and NTC cells. B) PARG protein expression levels relative to β-actin in HT29 PARG-KO and NTC (mean ± SD). The insert shows a representative image of a WB. C) Sensitivity of HT29 PARG-KO to FUIRI. Cells were treated with the indicated concentrations of 5FU + SN38 and stained with DiOC (viable cells) and DAPI (non-viable cells). The ratio of DAPI stained cells (non-viable cells) between HT29 PARG-KO and NTC was calculated for each treatment dilution. The arrow indicates the IC_50_ of the FUIRI treatment. Results are expressed as percentages. D) DePARylation activity and DNA damage levels in HT29 PARG-KO cells. Parental HT29, NTC, and PARG-KO cells were treated or not with 3.33 µM 5FU + 1.67 nM SN38 with or without 5 µM of PDD as indicated. PAR chain accumulation and DNA damage were detected by WB using antiPAR and H2A.X phosphorylated at Ser139 (γH2A.X) antibodies, respectively. The non-phosphorylated form of H2A.X protein was used as a normalizer. E) FUIRI-induced DNA damage in HT29 PARG-KO cells. Immunoflourescence staining of γH2A.X foci (red) shows increased DNA damage after 24h of FUIRI treatment in HT29 PARG-KO cells as compared to NTC cells. Nuclei were stained in blue (DAPI). Objective lens: 63X immersion oil. Scale bar, 30 µm. F) Sensitivity of HT29 PARG-KO cells to 5FU, SN38 and FUIRI. HT29 NTC and PARG-KO cells were treated for 24 hours with serial dilutions of 5FU, SN38, and FUIRI as indicated. Cells were stained with DiOC and DAPI, and cell viability (percentage of viable DiOC stained cells) was analyzed by flow cytometry. Graphs represent the mean percentage of dead HT29 PARG-KO cells relative to dead HT29 NTC cells (mean ± SD). G) Treatment-induced apoptosis in HT29 PARG-KO cells. HT29 NTC and PARG-KO cells were treated with different concentrations of 5FU, SN38 and FUIRI for 72 hours, as indicated. Apoptosis was analyzed by flow cytometry using the FITC Annexin V and propidium iodide staining. Bars represent the mean percentage of apoptotic cells ± SD. Differences in cell viability were analyzed using the Student’s t-test (*p < 0.05; **p < 0.01; ***p < 0.001).

To validate the functionality of our KO model, we investigated its dePARylation capacity following FUIRI treatment. A notable accumulation of PAR chains in HT29 PARG-KO cells compared to parental and HT29 NTC cells after 24 hours of treatment is illustrated in Figure 2D. When a PARG inhibitor (PDD) was added to FUIRI treatment to parental and HT29 NTC cells an increase in PAR chain accumulation was observed, indicating a deficiency in DNA dePARylation capacity. Given that defects in dePARylation are associated with increased DNA damage, we further analyzed the effect on H2A.X phosphorylation after FUIRI treatment using WB and immunofluorescence. Figures 2D and 2E show that, after 24 hours of FUIRI treatment, levels of γH2A.X were higher in HT29 PARG-KO cells as compared to HT29 or HT29-NTC cells, indicating an increase in cumulative DNA damage. This effect progressively increased over time (Supplementary Figure S6). These results suggest an impaired DNA repair mechanisms in our PARG-KO model. We then examined the impact of *PARG* deficiency on response to FUIRI as well as to each drug individually. Our findings, revealed that treatment with FUIRI led to a significant 33-70% increase in cell sensitivity to treatment in HT29 PARG-KO cells compared to the HT29 NTC line (Figure 2F). Notably, even at the lowest concentration of the drug combination, we observed a substantial 61% increased sensitivity with no further enhancement in effect at higher doses; efficacy remained consistent across the dosage spectrum. Treatment with 5FU exhibited an even more pronounced effect, whereas SN38 alone demonstrated a negligible impact on cell viability in the PARG-KO model as compared to NTC cells. These results aligned with those observed in apoptosis activation measurements (Figure 2G). Treatment with 5FU led to a greater increase in the percentage of apoptotic cells in PARG-KO cells across all tested doses. In contrast, no differences were observed with SN38 treatment at any dose. Therefore, the effect seen after combination treatment in PARG-KO cells is likely attributable to 5FU. These findings suggest that the genetic ablation of *PARG* in HT29 cells, compromises the repair of damage induced by the combination of 5FU and irinotecan, leading to heightened sensitivity and increased apoptosis. This effect appears to be primarily driven by the impact of 5FU

### 3.3. The Combination of PDD00017273 with 5FU, SN38, or Both Exhibits Synergistic Effects in Various CRC Cell Lines

Based on our results, we aimed to explore potential synergistic interactions between a pharmacological PARG inhibitor and the administration of 5FU and SN38, either separately or in combination. Among the available inhibitors (see Table 1), we selected the quinazolinedione-type PARG inhibitor PDD00017273 (PDD) for its high selectivity and efficiency in PARG inhibition, and because it had been extensively investigated in different solid tumor cell models (19–21). In the HT29 cell line, PDD exhibited no discernible impact on cell viability when administered as a standalone agent (Supplementary Figure S7). Therefore, for subsequent synergism evaluations, we selected 1 μM and 5 μM doses based on their demonstrated PARG inhibitory effects in prior investigations (19–21). As shown in Figure 3, treatment with the lowest dose of PDD sensitized HT29 cells to 5FU, SN38 and their combination. Notably, this sensitization effect was substantially potentiated and displayed pronounced synergy when PDD was administered at 5 μM. Further investigations extended to the HT115 and DLD1 cell lines revealed both similarities and distinctions. In HT115 cells, PDD conferred sensitivity to SN38 and FUIRI treatment only when it was added at 5 µM, whereas sensitivity to 5FU was enhanced at both concentrations tested. However, in the DLD1 cell line, although the effect of PDD on SN38 and FUIRI is similar to that observed in the other cell lines, only the highest concentration of PDD enhanced the efficacy of 5FU treatment. Synergism studies using the Chou and Talalay method revealed that the combination of PDD at the 5 µM concentration was synergistic in all cases and cell lines. However, when used at 1 µM, it was additive or slightly antagonistic in a few cases (Supplementary Figure S8). These results indicate that pharmacological inhibition of PARG acts synergistically with 5-FU, SN38, and their combination, though the strength of this effect varies depending on the context.

**Figure 3.**
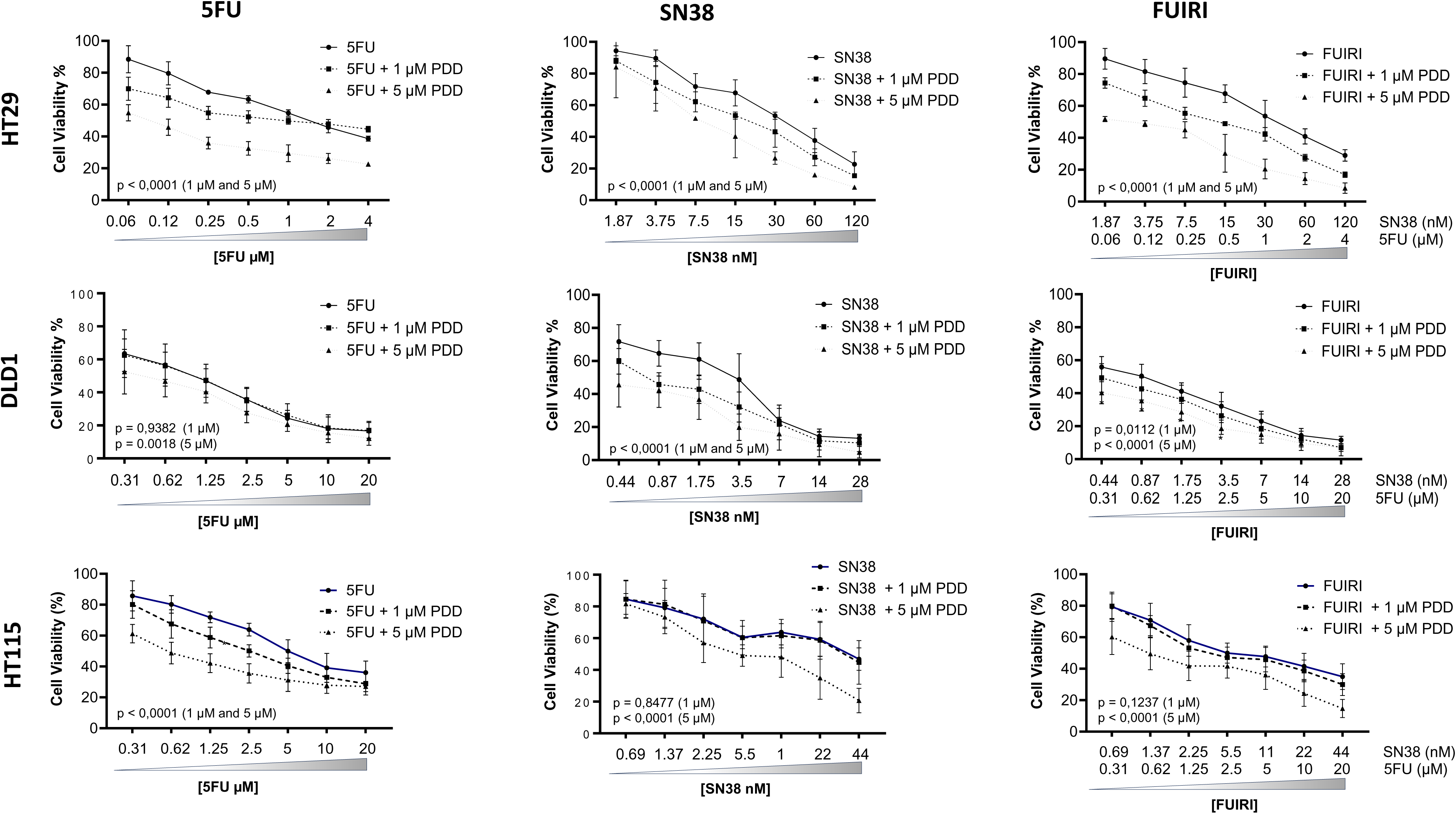
Effect of the combination of PDD with 5FU, SN38 and FUIRI on CRC cell lines. HT29, DLD1 and HT115 cells were treated with different concentrations of 5FU, SN38 and FUIRI as indicated, with or without the addition of 1 µM or 5 µM of PDD for 72 hours. Cell viability was assessed using the MTT assay. Differences between treatments were analysed using a two-way ANOVA test. The p-values shown correspond to the comparison between combination treatment with PDD and chemotherapy alone. Graphs represent the mean percentage of cell viability ± SD.

### 3.4. PARG-KO Tumors Exhibit Increased Sensitivity to 5FU Treatment *In Vivo*

The low bioavailability of PDD makes it unsuitable for *in vivo* applications. However, at the time of this study, another inhibitor, COH34, showed promise, particularly in *PARP*-mutated ovarian cancer models. We treated HT29 cells with various concentrations of COH34, both alone and in combination with 5FU, SN38, or FUIRI. Our results (Supplementary Figure S9) indicated that COH34 was ineffective in all conditions. Therefore, to investigate the potential synergism between PARG inhibition and 5FU or FUIRI treatment *in vivo*, we subcutaneously injected HT29 PARG-KO or NTC cells into immunodeficient mice and treated them with these drugs (Figure 4A). Tumor growth was monitored for 21 days in the absence of any treatment or drug vehicle, revealing no significant differences in relative tumor volume between both groups (Figure 4B). This observation aligns with our *in vitro* results, which showed no effect of PARG-KO on cell proliferation. For the subsequent experiments, the doses of 5FU (50 mg/kg) and irinotecan (50 mg/kg), as well as the administration schedule (once a week for 4 weeks), were selected based on previous studies, the experience of our close collaborators and proving to be non-toxic to animals (Supplementary Figure S10). Notably, these doses are lower than those typically administered to patients (22). Both treatments reduced the growth rate (RTV) of HT29 PARG-KO and NTC tumors with a more pronounced effect observed in the HT29 PARG-KO tumours (Figure 4C). The tumour growth inhibition rate (TGI) indicated that 5FU had a slightly stronger effect in the absence of *PARG* expression, while there was no significant difference in the effect of FUIRI treatment (Figure 4D). While these results are not definitive, they are consistent with our *in vitro* findings and suggest that at least, the combination of 5FU and a PARG inhibitor may be beneficial for CRC patients.

**Figure 4.**
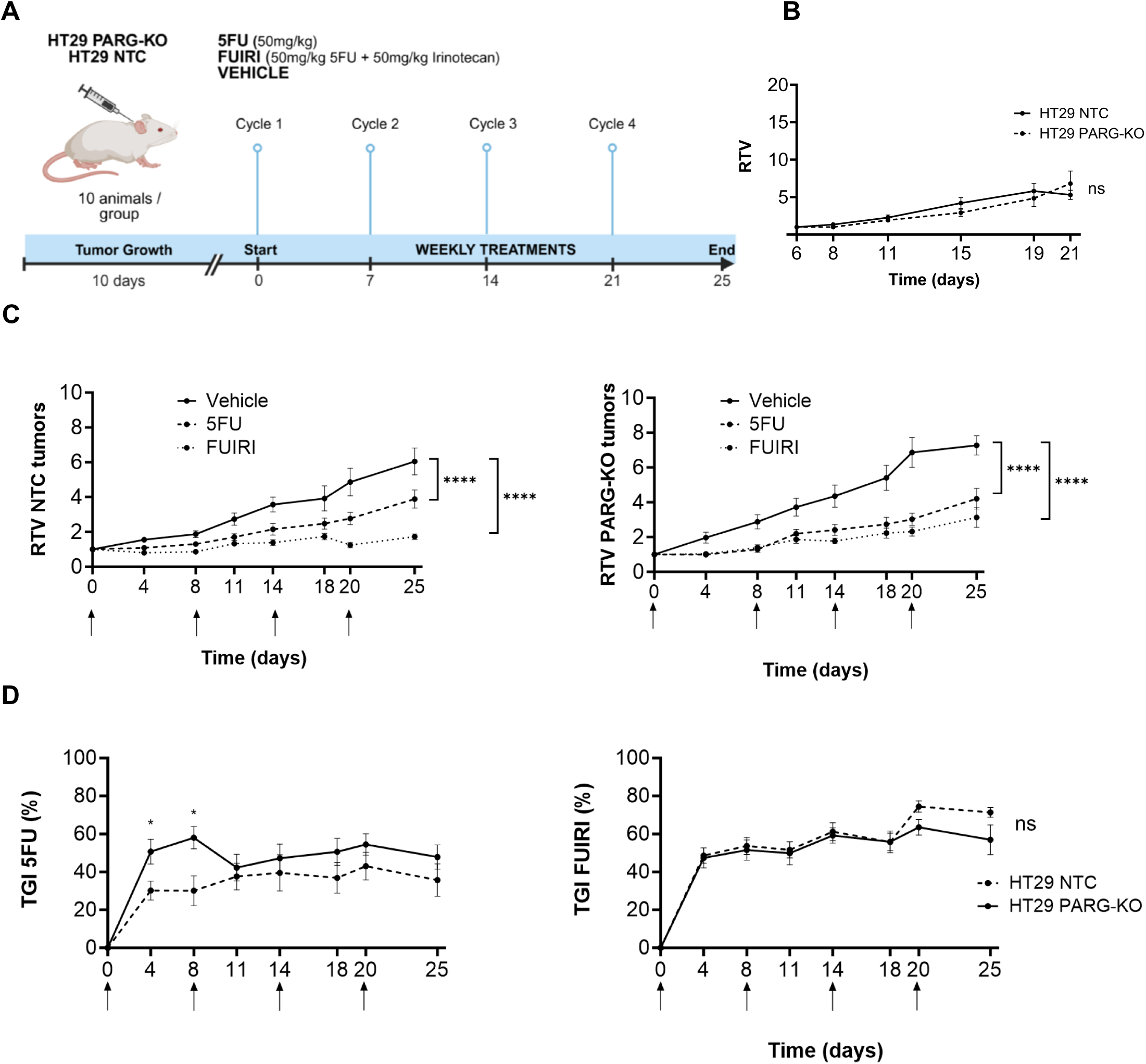
*In vivo* experiments. A) Study design. HT29 NTC and PARG-KO cells were subcutaneously injected in Balb/c mice and allowed to grow for 10 days before treatment was initiated. Tumour-bearing mice were treated with vehicle (PBS), 5FU or FUIRI at the indicated doses (10 mice per group). Treatments were administered intraperitoneally once per week. Mice were euthanized on day 25. B) Tumour growth. The volume of HT29 NTC and PARG-KO subcutaneous tumours was monitored twice per week for 21 days. Measurements prior to day 6 were not possible due to the low tumour volume. The graphs represent the mean Relative Tumour Volume (RTV: ratio between the tumour volume on day *x* and the tumour volume on day *0*) ± SEM. C) Effect of 5FU or FUIRI treatment on mice bearing HT29 PARG-KO or NTC tumours. Tumour size was measured twice per week until the end of the experiment. Graphs show mean RTV ± SEM for the indicated treatments. D) Tumour Growth Inhibition Rate (TGI). The percentage of TGI induced by 5FU or FUIRI treatments in mice bearing HT29 PARG-KO and NTC tumours, was calculated using the formula (1 - RTV) * 100. Arrows indicate the days of treatment administration for 5FU and FUIRI. Statistical significance in panels B, C, and D were determined using a Linear Mixed Effects (LME) model followed by a post hoc Estimated Marginal Means (EMM) analysis (*p < 0.05).

### 3.5. Investigating PARG Expression as a Predictive Biomarker for 5FU + Irinotecan Treatment in Metastatic CRC patients

Finally, we aimed to investigate whether tumour expression of PARG could serve as a predictive biomarker for treatment with 5FU + irinotecan. We utilized a retrospective cohort of 170 primary FFPE tumour samples from metastatic CRC patients treated with first-line irinotecan + 5FU and assessed PARG protein expression via IHC. Patients’ characteristics can be seen in Table 2. HT29 PARG-KO and NTC cells were used as negative and positive staining controls, respectively (Figure 5A) and helped us establish the positivity threshold in CRC primary tumor samples. PARG staining of tumours was classified as null, slight (1+), medium (2+) or strong (3+). We found a high percentage of samples (61%) being negative for PARG expression (Figure 5B).

**Table 2.** Patients’ characteristics

| Characteristic |  | PARG null | PARG other | p-value | All |
| --- | --- | --- | --- | --- | --- |
| Sex, N (%) | Woman | 28 (62) | 17 (38) | 0.58 | 45 (26.5) |
|  | Man | 65 (67) | 32 (33) |  | 97 (57) |
|  | N/A |  |  |  | 28 (16.5) |
| Age, mean [range] |  | 61.65 | 61.59 | 0.79 | 61.63 |
| Localization N (%) | Colon | 83 (65.4) | 44 (34.6) | 0.35 | 127 (75) |
|  | Rectum | 28 (70) | 12 (30) |  | 40 (24) |
|  | Both | 2 (100) | 0 (0) |  | 2 (1) |
| ECOG N (%) | 0 | 60 (66.7) | 30 (33.3) | 0.87 | 90 (53) |
|  | 1 | 46 (66.7) | 23 (33.3) |  | 69 (40.5) |
|  | 2 | 4 (57.1) | 3 (42.9) |  | 7 (4) |
|  | N/A |  |  |  | 9 (2.5) |
| Metastases N (%) | Liver only | 54 (66.7) | 27 (33.3) | 0.71 | 81 (49) |
|  | Lung only | 16 (72.7) | 6 (27.3) |  | 22 (13) |
|  | Other | 4 (80) | 1 (20) |  | 5 (3) |
|  | Multiple | 37 (61.7) | 23 (38.4) |  | 60 (22) |
| 1st line N (%) | FUIRI | 58 (62.5) | 31 (34.8) | 0.75 | 89 (52) |
|  | FOLFIRI | 55 (67.9) | 26 (32.1) |  | 81 (48) |
| Response N (%) | R | 71 (70.3) | 30 (29.7) | 0.115 | 101 (59.4) |
|  | NR | 32 (57.1) | 24 (42.9) |  | 56 (33) |
|  | N/A |  |  |  | 13 (7.6) |
| TTP, median [95% CI] |  | 8.95 [7.75 – 10.15] | 8.42 [8.02 – 8.82] | 0.55 | 8.59 [7.98 – 9.29] |
| OS, median [95% CI] |  | 24.18 [18.23 – 30.06] | 22.66 [20.12 – 25.21] | 0.21 | 22.66 [19.92 – 25.41] |
| OS>20 months |  | 36.45 [26.94 – 45.96] | 27.90 [23.81 – 31.98] | 0.01 | N/A |

PARG positive = PARG IHC staining was +1, +2 or +3; FUIRI = irinotecan (80 mg/m<sup>2</sup>) + 5FU (250 mg/m<sup>2</sup> 48-h infusion) weekly; FOLFIRI = irinotecan (180 mg/m<sup>2</sup> on day 1) + 5FU (400 mg/m<sup>2</sup> bolus and 600 mg/m<sup>2</sup> 22-h infusion) + leucovorin (LV) (200 mg/m<sup>2</sup> on days 1-2) every 2 weeks; R = Responders (complete + partial response); NR = non-responders (stable + progressive disease); TTP = time to progression; OS = overall survival; CI = confidence

**Figure 5.**
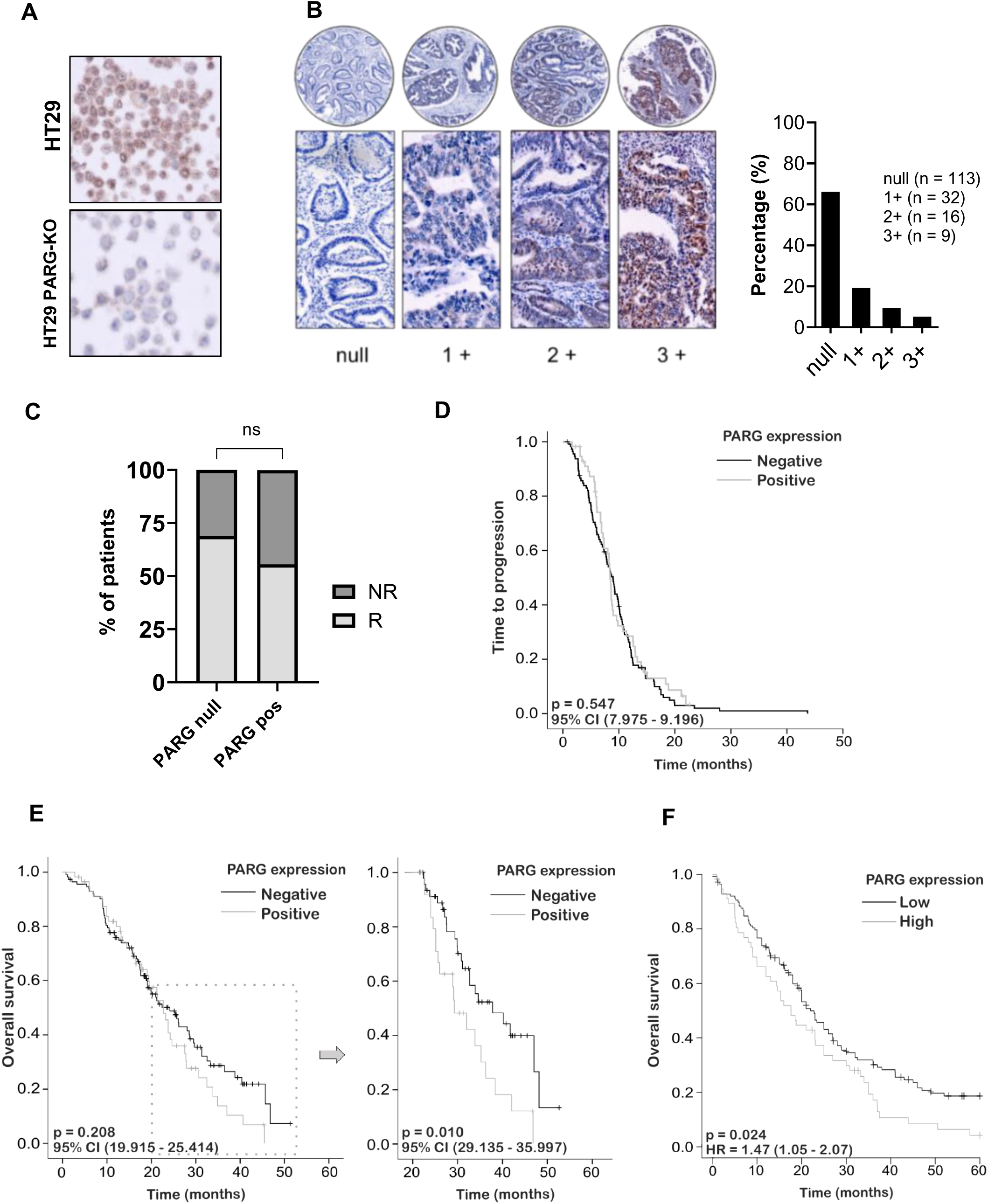
PARG expression and its association with clinical outcomes of CRC patients treated with 5FU + irinotecan. A) PARG IHC staining. After embedding in paraffin, HT29 PARG-KO and NTC were used as negative and positive controls for PARG IHC staining, respectively. B) PARG IHC staining of FFPE CRC primary tumours. Representative images of CRC tissue sections showing different degrees of PARG IHC staining (left). Percentage and total number of samples in each staining category (right). C) Response to treatment according to PARG IHC staining. Patients were categorized based on PARG IHC staining, as null or positive (1+, 2+ or 3+). The graph shows the percentage of patients responding (partial or complete response) or non-responding (stable or progressive disease) in each of these two categories. Differences between categories were analysed using Fisher’s exact test. D) Time to progression (TTP) of CRC patients according to PARG IHC staining. Kaplan-Meier plot shows the TTP probability for patients with null or positive PARG IHC staining. E) Overall survival (OS) of CRC patients according to PARG IHC staining. Kaplan-Meier plot shows OS probability for patients with null or positive PARG IHC staining (left). Kaplan-Meier plot representing OS beyond 20 months according to PARG IHC staining (right). F) OS of stage IV CRC patients according to PARG mRNA expression levels as reported by the the KM plotter platform. Patients were classified as having low or high PARG mRNA expression using the best cutoff method. In panels D, E, and F, differences between groups were calculated using the log rank test. CI = Confidence Interval; HR = Hazard ratio.

Therefore, to more effectively categorize the samples into homogeneous groups, we contrasted the negative cases with those exhibiting either positivity. Nearly 70% of the PARG-negative patients responded to the treatment, whereas this percentage decreased to 56% in the positive cases, showing a trend toward statistical significance (Figure 5C) (*X*^2^ p = 0.096; Fisher’s test p=0.115; Table 2). PARG has been associated with increased metastatic potential (23) and indeed, we observed that patients with null PARG expression had a significantly lower number of liver metastases (≤3 metastases) compared with those expressing PARG (Supplementary Figure S11). There was no difference in TTP (Figure 5D) or OS up to 20 months (Figure 5E; left) according to negative or positive PARG expression. However, a significantly higher percentage of PARG-negative patients survived beyond 20 months compared to PARG-positive patients (Figure 5E; right). Lastly, using the online application Kapplan Meier Plotter (24) we observed that low *PARG* gene expression (data from GEO datasets) in stage IV CRC tumours was associated with improved OS in patients (Figure 5F). Unfortunately, data on the first-line treatments received by these patients are unavailable. These findings suggest that the absence of PARG expression in primary tumours of mCRC patients could be indicative of a better long-term clinical outcome when treated with combination regimens of 5FU and irinotecan. However, further research is needed to confirm its value as predictive biomarker.

## 4. Discussion

CRC is a globally prevalent disease, with chemotherapy achieving response rates of approximately 50% in metastatic cases. However, resistance to treatment often leads to disease progression, underscoring the need for new therapeutic strategies. To address this challenge, researchers are investigating chromatin regulator genes, which modulate DNA accessibility and influence treatment response, as potential drug targets.

Through a high-throughput LOF screen, we identified PARG as a key regulator of CRC treatment sensitivity. PARG plays a crucial role in DNA repair by degrading PAR chains. Our findings demonstrate for the first time, that inhibiting PARG, either genetically or pharmacologically, enhances sensitivity to chemotherapy, particularly to 5FU and the FUIRI combination and suggest that PARG inhibition could be a promising strategy to improve treatment efficacy in CRC patients.

Our approach leveraged the hEPI9 shRNA library, which targets over 900 chromatin regulator genes and has been successfully used in previous LOF screenings by our group (6,25). Among the top hits, we identified *ATR*, a serine/threonine protein kinase that activates checkpoint signaling in response to genotoxic stress, promoting DNA damage repair. Notably, several studies have highlighted the synergistic effects of ATR inhibitors with 5FU and irinotecan (12–14). Other key candidates included *ATM*, *BRCA1*, which are involved in the HR repair pathway and play crucial roles in repairing DNA double-strand breaks (26–28). Additionally, we identified *PARP14* and *PARG*, whose depletion leads to DNA damage accumulation (29,30). A Gene Ontology analysis further reinforced strong associations between our candidate genes and DNA repair processes. Mechanistically, 5FU treatment inhibits TS, resulting in a disrupted nucleotide pool and the misincorporation of 5-FdUTP or d-UTP into DNA. This misincorporation leads to increased DNA damage, including single- and double-strand breaks (31,32). Meanwhile, SN38 inhibits topoisomerase 1, preventing the re-ligation of DNA strands and further facilitating double-strand break formation (33). Taken together, these findings validate the robustness and reliability of our screening approach, providing key insights into potential therapeutic targets for CRC treatment.

We selected PARG as the focus of our study due to the availability of pharmacological inhibitors already in clinical trials and the limited information on its role in CRC. PARG functions as a glycohydrolase enzyme, responsible for degrading PAR chains, a process with two critical effects: (1) Mono-ADP-ribose moieties are released and metabolized into ATP, supporting essential metabolic processes and signaling pathways; (2) Longer PAR fragments (more than three ADP units) act as apoptotic signals, influencing cell fate (29,34). As the primary regulator of protein PARylation, PARG plays a central role in the DNA damage response (DDR), particularly in coordination with PARP1. While other hydrolases—such as TARG1, ARH1, ARH3, MacroD1, and MacroD2— contribute to this process, their impact is relatively minor (35).

Upon DNA damage induced by chemotherapeutic agents, PARP1 recognizes single- or double-strand breaks, initiating PARylation and recruiting DNA damage response proteins to start the repair process. PARG subsequently dePARylates these factors, allowing their redistribution to new DNA damage sites (34).

Therefore, PARG inhibition disrupts this repair cycle, leading to PAR chain accumulation, increased DNA damage, and impaired repair mechanisms (36). Indeed, we observed a significant accumulation of PAR chains and increased DNA damage in HT29 cells treated with FUIRI, whereas these effects were absent in untreated cells. Notably, these effects were even more pronounced in HT29 PARG-KO cells or when treatment was combined with the PDD inhibitor. Our findings align with previous studies demonstrating that PARG depletion enhances the efficacy of chemotherapeutic agents in melanoma, ovarian cancer, glioblastoma, and head and neck cancer (37–40). Moreover, the literature supports a connection between increased treatment sensitivity and apoptosis, suggesting that PAR chain accumulation—due to reduced PARG activity—along with DNA damage accumulation from impaired repair mechanisms, triggers the release of apoptotic-inducing factor (AIF) from mitochondria, ultimately driving apoptosis (29).

Our data indicate that PARG inhibition enhances the sensitivity of CRC cell lines to 5FU and SN38, though the effects differ between genetic knockout and pharmacological inhibition. Thus, in HT29 PARG-KO cells, sensitivity to 5FU and 5FU + SN38 was increased (reduced viability and increased apoptosis) but sensitivity to SN38 alone remained largely unaffected. In contrast, treatment with the PDD increased sensitivity to 5FU, SN38, and 5FU + SN38 across HT29, HCT115, and DLD1 cell lines.

A key factor influencing this difference may be the presence of gain-of-function (GOF) *TP53* mutations (R273H in HT29, S241F in HCT15 and DLD1), which are known to promote chemoresistance and adaptation to replication stress (41). In PARG-KO HT29 cells, mutant TP53 may activate compensatory repair mechanisms, such as alternative pathways for resolving TOP1-DNA-Protein Crosslinks (DPCs), reducing the expected sensitivity to SN38. Conversely, acute pharmacological PARG inhibition prevents such adaptation, leading to persistent PARylation accumulation and impaired TOP1-DPC degradation, thereby enhancing SN38 cytotoxicity (42).

These findings were further validated *in vivo*, where HT29 parental and PARG-KO cells were subcutaneously implanted in athymic mice and treated with 5FU or 5FU + SN38. The results showed that 5FU treatment was more effective in reducing tumor growth in PARG-KO tumors than in parental tumors while 5FU + SN38 treatment showed no significant differences between the two groups, suggesting the activation of compensatory mechanisms under these conditions. However, some limitations must be considered here, such as the use of subcutaneous instead of orthotopic tumors and the reliance on a KO model rather than a pharmacological inhibitor.

This is the first study to suggest a potential synergy between PARG inhibitors, 5FU, and possibly SN38—either alone or in combination—in CRC. While no previous studies have explored the pharmacological effect of combining PARG and topoisomerase 1 inhibitors in cancer, two studies led by Dr. Jonathan R. Brody have reported a synergistic effect between PARG inhibitors and 5FU or CF10 (a next-generation fluoropyrimidine) in pancreatic ductal adenocarcinoma (43,44). Nearly all CRC patients receive 5FU at some point in their treatment, whether in the adjuvant setting or for advanced disease. Every therapeutic combination includes this fluoropyrimidine or one of its derivatives, such as capecitabine. Even in patients who develop resistance to oxaliplatin- or irinotecan-based regimens, 5FU remains a cornerstone of subsequent treatment lines. Therefore, our findings suggest that combining 5FU with a PARG inhibitor could have a significant clinical impact, offering a novel therapeutic strategy that enhances existing treatment options in CRC.

Finally, we explored the potential of PARG as a predictive biomarker for first-line treatment with 5FU and irinotecan-based combinations. Our results suggest a possible association between PARG positivity (assessed by IHC) and lower response rates as well as reduced long-term clinical benefit. However, the lack of correlation with time to progression makes these findings inconclusive.

A key limitation here is that tumor samples were obtained from primary tumors at diagnosis, rather than from the metastases that were actually treated. As a result, we cannot determine whether PARG expression remained stable or changed during tumor progression. It is possible that PARG expression increases during metastasis, meaning that patients initially classified as PARG-negative may have actually developed PARG positivity over time. This could explain the lack of significant differences between groups. In line with this, the work of Wang JQ et al, demonstrated that PARG silencing in CT26 mouse CRC cells compromised metastasis formation when these cells were injected into the spleen of immunocompetent mice (45). Furthermore, our own results indicate that 78% of patients with PARG-negative tumors had 3 or fewer liver metastases, while this percentage was lower (58%) in those with PARG expression (Fisher’s test p = 0.01). Nevertheless, further studies in larger cohorts using metastatic samples are needed to validate this hypothesis.

Notably, our findings showing a long-term benefit in patients with PARG-negative tumors, were replicated in an independent cohort. Additionally, similar associations have been reported in other cancer types, including hepatocellular carcinoma and breast cancer (46,47).

Currently, two PARG inhibitors are in clinical trials. ETX-19477 (858 Therapeutics) entered a Phase I trial in May 2024 to evaluate safety, pharmacokinetics (PK), pharmacodynamics (PD), and antitumor activity. Preclinical studies demonstrated inhibition of proliferation and increased γH2AX levels across multiple cancer cell lines, with *in vivo* efficacy in breast and ovarian xenograft models (17,48). IDE-161 (IDEAYA) initiated a Phase I trial in April 2023, investigating PARG inhibition in combination with immunotherapy. The study assesses the safety, tolerability, PK, PD, and antitumor activity of IDE-161 alone or with pembrolizumab (anti-PD-1) in patients with BRCA1/2 mutations or HR defects. Additionally, IDEAYA is planning a new Phase I trial combining IDE-161 with pembrolizumab for patients with MSI-high and MSS endometrial cancer (18,49).

In conclusion, our findings open a promising avenue for clinical research, potentially marking a significant advance in the treatment of metastatic CRC. The absence of new first-line approvals for over two decades—largely due to the failure of immune checkpoint inhibitors in MSS patients and PARP1 inhibitors—underscores the urgent need for novel therapeutic targets. Our results could contribute meaningfully to enhancing both survival and quality of life for patients with this challenging malignancy.

## Acknowledgements

This work was been supported by the FEDER/Ministerio de Ciencia e Innovación - Agencia Estatal de Investigación through the grant PIE16/00011 RESPONSE to Dr. Marcus Buschbeck, by the ISCIII grant from the Spanish Government, project number PI20/01183 and the Departament d’Innovació, Universitats i Empresa, Generalitat de Catalunya, project number 2017-SGR-723, both of them awarded to Dr. Martinez-Balibrea, Ministerio de Ciencia e innovacion grant PID2019-105278RB-I00 and Fundación Olaga Torres Josep Bombardó Navinés Award both Dr. Sonia Forcales. Ferran Grau Leal is supported by an AGAUR-FI grant (2023 FI-3 00065) from the predoctoral program Joan Oró of the Secretaria d′Universitats i Recerca del Departament de Recerca i Universitats de la Generalitat de Catalunya and the European Social Funds Plus. Carla Vendrell Ayats holds an INVESTIGO program contract from the Generalitat de Catalunya, funded by the EU, Next Generation European Funds. Jeannine Diesch is supported by a grant from the Scientific Foundation of the Spanish Association Against Cancer (INVES223200DIES). Research in the Buschbeck lab is further supported by the following grants: the Marie Skłodowska Curie Doctoral Network ‘NUCLEAR’ (HORIZON-MSCA-2023-DN-101166838), LaCaixa Banking Foundation (CI24-10180), AGAUR 2021-SGR-260, and PRYGN222668BUSC from the Fundación AECC. This work was also funded by “Bates Blanques” solidary movement (https://www.lesbatesblanques.cat/).

The authors declare there is no conflict of interest. All authors have read the journal’s authorship agreement and policy on disclosure of potential conflicts of interest.

## 6. Declaration of Generative AI and AI-assisted technologies in the writing process

During the preparation of this work, the authors used ChatGPT 3.5 in order to improve readability and language. After using this tool, the authors reviewed and edited the content as needed and take full responsibility for the content of the publication.

---

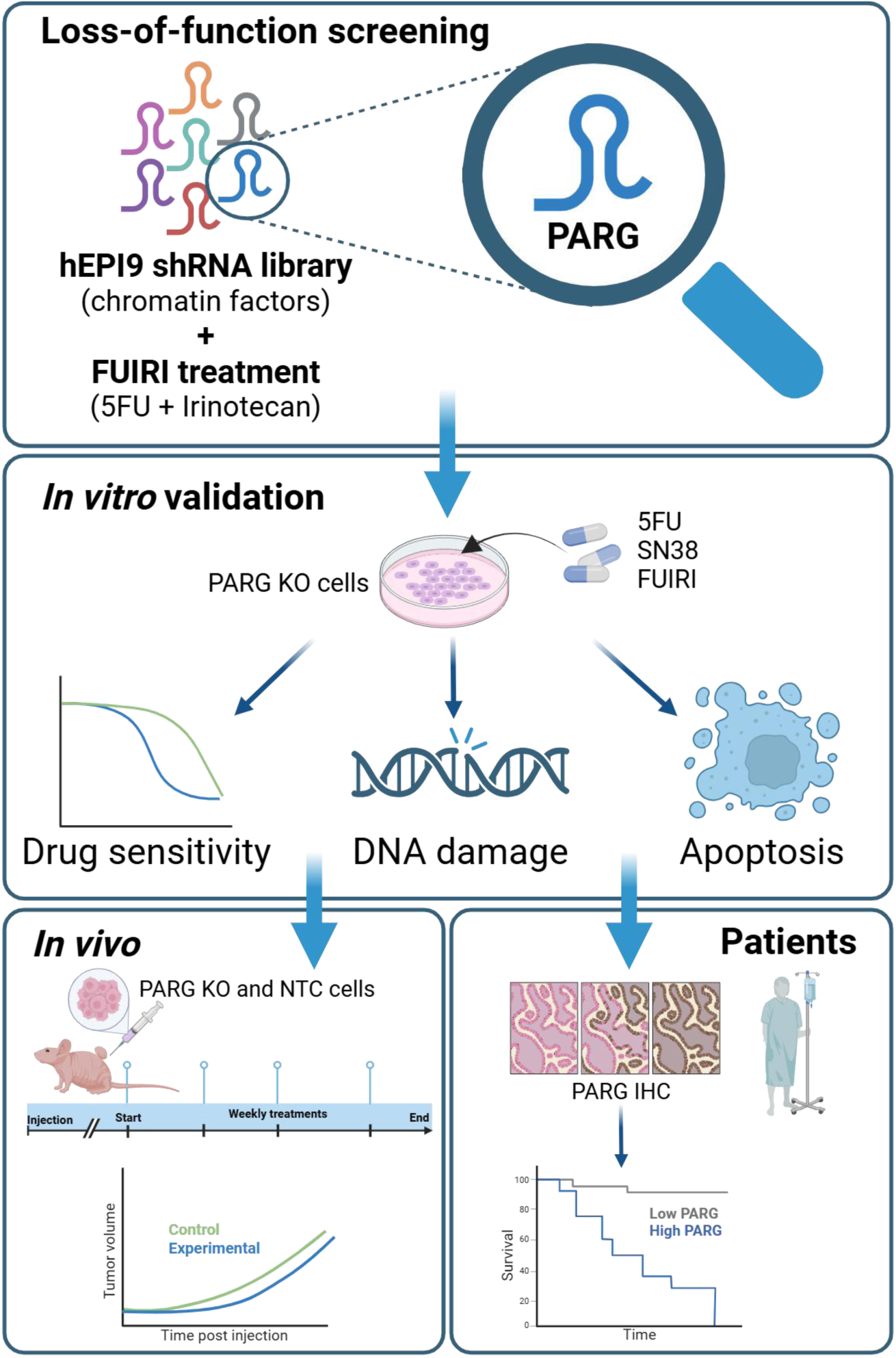

## Bibliography

1. Saltz LB, Cox J V., Blanke C, Rosen LS, Fehrenbacher L, Moore MJ, et al. Irinotecan plus fluorouracil and leucovorin for metastatic colorectal cancer. Irinotecan Study Group. N Engl J Med [Internet]. 2000 Sep 28 [cited 2025 Feb 14];343(13):905–14. Available from: https://pubmed.ncbi.nlm.nih.gov/11006366/

2. Moreta-Moraleda C, Queralt C, Vendrell-Ayats C, Forcales S, Martínez-Balibrea E. Chromatin factors: Ready to roll as biomarkers in metastatic colorectal cancer? Pharmacol Res. 2023 Oct 1;196:106924.

3. Ritchie ME, Dai Z, Sheridan JM, Gearing LJ, Moore DL, Su S, et al. edgeR: a versatile tool for the analysis of shRNA-seq and CRISPR-Cas9 genetic screens. F1000Res [Internet]. 2014 [cited 2024 Sep 9];3. Available from: /pmc/articles/PMC4023662/

4. Aranda E, Valladares M, Martinez-Villacampa M, Benavides M, Gomez A, Massutti B, et al. Randomized study of weekly irinotecan plus high-dose 5-fluorouracil (FUIRI) versus biweekly irinotecan plus 5-fluorouracil/leucovorin (FOLFIRI) as first-line chemotherapy for patients with metastatic colorectal cancer: A Spanish Cooperative Group for the Treatment of Digestive Tumors Study. Annals of Oncology [Internet]. 2009 Feb 1 [cited 2024 Sep 9];20(2):251–7. Available from: http://www.annalsofoncology.org/article/S0923753419404717/fulltext

5. Fellmann C, Zuber J, McJunkin K, Chang K, Malone CD, Dickins RA, et al. Functional Identification of Optimized RNAi Triggers Using a Massively Parallel Sensor Assay. Mol Cell [Internet]. 2011 Mar 18 [cited 2024 Sep 10];41(6):733–46. Available from: http://www.cell.com/article/S1097276511000918/fulltext

6. Fellmann C, Hoffmann T, Sridhar V, Hopfgartner B, Muhar M, Roth M, et al. An optimized microRNA backbone for effective single-copy RNAi. Cell Rep [Internet]. 2013 Dec 26 [cited 2024 Sep 9];5(6):1704–13. Available from: http://www.cell.com/article/S2211124713006876/fulltext

7. Martinez-Balibrea E, Plasencia C, Ginés A, Martinez-Cardús A, Musulén E, Aguilera R, et al. A proteomic approach links decreased pyruvate kinase M2 expression to oxaliplatin resistance in patients with colorectal cancer and in human cell lines. Mol Cancer Ther [Internet]. 2009 Apr 1 [cited 2024 Sep 10];8(4):771–8. Available from: /mct/article/8/4/771/93394/A-proteomic-approach-links-decreased-pyruvate

8. Martinez-Cardús A, Martinez-Balibrea E, Bandrés E, Malumbres R, Ginés A, Manzano JL, et al. Pharmacogenomic approach for the identification of novel determinants of acquired resistance to oxaliplatin in colorectal cancer. Mol Cancer Ther [Internet]. 2009 Jan 1 [cited 2024 Sep 10];8(1):194–202. Available from: /mct/article/8/1/194/93185/Pharmacogenomic-approach-for-the-identification-of

9. Ruiz De Porras V, Bystrup S, Martínez-Cardús A, Pluvinet R, Sumoy L, Howells L, et al. Curcumin mediates oxaliplatin-acquired resistance reversion in colorectal cancer cell lines through modulation of CXC-Chemokine/NF-κB signalling pathway. Sci Rep [Internet]. 2016 Apr 19 [cited 2024 Sep 10];6. Available from: https://pubmed.ncbi.nlm.nih.gov/27091625/

10. Sveen A, Bruun J, Eide PW, Eilertsen IA, Ramirez L, Murumagi A, et al. Colorectal cancer consensus molecular subtypes translated to preclinical models uncover potentially targetable cancer cell dependencies. Clinical Cancer Research [Internet]. 2018 Feb 15 [cited 2024 Sep 10];24(4):794– 806. Available from: /clincancerres/article/24/4/794/81198/Colorectal-Cancer-Consensus-Molecular-Subtypes

11. Berg KCG, Eide PW, Eilertsen IA, Johannessen B, Bruun J, Danielsen SA, et al. Multi-omics of 34 colorectal cancer cell lines - a resource for biomedical studies. Mol Cancer [Internet]. 2017 Jul 6 [cited 2024 Sep 10];16(1):1–16. Available from: https://molecular-cancer.biomedcentral.com/articles/10.1186/s12943-017-0691-y

12. Jossé R, Martin SE, Guha R, Ormanoglu P, Pfister TD, Reaper PM, et al. ATR inhibitors VE-821 and VX-970 sensitize cancer cells to topoisomerase I inhibitors by disabling DNA replication initiation and fork elongation responses. Cancer Res [Internet]. 2014 Dec 1 [cited 2025 Feb 14];74(23):6968. Available from: https://pmc.ncbi.nlm.nih.gov/articles/PMC4252598/

13. Villaruz LC, Kelly K, Waqar SN, Davis EJ, Shapiro G, LoRusso P, et al. NCI 9938: Phase I clinical trial of ATR inhibitor berzosertib (M6620, VX-970) in combination with irinotecan in patients with advanced solid tumors. Journal of Clinical Oncology [Internet]. 2022 Jun 1 [cited 2025 Feb 14];40(16_suppl):3012–3012. Available from: https://ascopubs.org/doi/10.1200/JCO.2022.40.16_suppl.3012

14. Clinical Trial: NCT04535401 - My Cancer Genome [Internet]. [cited 2025 Feb 14]. Available from: https://www.mycancergenome.org/content/clinical_trials/NCT04535401/

15. Study Details | A Study of PARG Inhibitor IDE161 in Participants With Advanced Solid Tumors | ClinicalTrials.gov [Internet]. [cited 2024 Oct 7]. Available from: https://clinicaltrials.gov/study/NCT05787587

16. Study Details | A Study of PARG Inhibitor ETX-19477 in Patients With Advanced Solid Malignancies | ClinicalTrials.gov [Internet]. [cited 2024 Oct 7]. Available from: https://clinicaltrials.gov/study/NCT06395519

17. Holleran JP, Rodems TS, Sharma S, Santini AM, Wuerz C, Liu J, et al. Abstract 2083: Discovery of ETX-19477, a novel and selective PARG inhibitor with high potency against tumors with underlying replication stress. Cancer Res [Internet]. 2024 Mar 15 [cited 2024 Oct 7];84(6_Supplement):2083–2083. Available from: /cancerres/article/84/6_Supplement/2083/736581/Abstract-2083- Discovery-of-ETX-19477-a-novel-and

18. Abed M, Muñoz D, Seshadri V, Federowicz S, Rao AA, Bhupathi D, et al. Abstract 6093: IDE161, a potential first-in-class clinical candidate PARG inhibitor, selectively targets homologous-recombination-deficient and PARP inhibitor resistant breast and ovarian tumors. Cancer Res [Internet]. 2023 Apr 1 [cited 2024 Oct 7];83(7_Supplement):6093–6093. Available from: /cancerres/article/83/7_Supplement/6093/723258/Abstract-6093- IDE161-a-potential-first-in-class

19. Gravells P, Neale J, Grant E, Nathubhai A, Smith KM, James DI, et al. Radiosensitization with an inhibitor of poly(ADP-ribose) glycohydrolase: A comparison with the PARP1/2/3 inhibitor olaparib. DNA Repair (Amst) [Internet]. 2018 Jan 1 [cited 2024 Oct 2];61:25–36. Available from: https://pubmed.ncbi.nlm.nih.gov/29179156/

20. Tsuda K, Kurasaka C, Ogino Y, Sato A. Genomic and biological aspects of resistance to selective poly(ADP-ribose) glycohydrolase inhibitor PDD00017273 in human colorectal cancer cells. Cancer Rep (Hoboken) [Internet]. 2023 Feb 1 [cited 2024 Oct 2];6(2). Available from: https://pubmed.ncbi.nlm.nih.gov/36053937/

21. James DI, Smith KM, Jordan AM, Fairweather EE, Griffiths LA, Hamilton NS, et al. First-in-Class Chemical Probes against Poly(ADP-ribose) Glycohydrolase (PARG) Inhibit DNA Repair with Differential Pharmacology to Olaparib. ACS Chem Biol [Internet]. 2016 Nov 18 [cited 2024 Oct 2];11(11):3179–90. Available from: https://pubmed.ncbi.nlm.nih.gov/27689388/

22. Nair AB, Jacob S. A simple practice guide for dose conversion between animals and human. J Basic Clin Pharm [Internet]. 2016 [cited 2024 Oct 3];7(2):27. Available from: /pmc/articles/PMC4804402/

23. Li Q, Li M, Wang YL, Fauzee NJS, Yang Y, Pan J, et al. RNA interference of PARG could inhibit the metastatic potency of colon carcinoma cells via PI3-kinase/Akt pathway. Cell Physiol Biochem [Internet]. 2012 [cited 2025 Mar 18];29(3–4):361–72. Available from: https://pubmed.ncbi.nlm.nih.gov/22508044/

24. Győrffy B. Integrated analysis of public datasets for the discovery and validation of survival-associated genes in solid tumors. Innovation (Cambridge (Mass)) [Internet]. 2024 May 6 [cited 2025 Mar 3];5(3). Available from: https://pubmed.ncbi.nlm.nih.gov/38706955/

25. Diesch J, Le Pannérer MM, Winkler R, Casquero R, Muhar M, van der Garde M, et al. Inhibition of CBP synergizes with the RNA-dependent mechanisms of Azacitidine by limiting protein synthesis. Nat Commun. 2021 Dec 1;12(1).

26. Jin MH, Oh DY. ATM in DNA repair in cancer. Pharmacol Ther [Internet]. 2019 Nov 1 [cited 2025 Feb 14];203. Available from: https://pubmed.ncbi.nlm.nih.gov/31299316/

27. Rosen EM. BRCA1 in the DNA damage response and at telomeres. Front Genet [Internet]. 2013 Jun 21 [cited 2025 Feb 14];4(JUN):52895. Available from: www.frontiersin.org

28. Katheeja MN, Das SP, Das R, Laha S. BRCA1 interactors, RAD50 and BRIP1, as prognostic markers for triple-negative breast cancer severity. Front Genet. 2023;14.

29. Feng X, Koh DW. Roles of poly(ADP-ribose) glycohydrolase in DNA damage and apoptosis. Int Rev Cell Mol Biol [Internet]. 2013 [cited 2025 Feb 14];304:227–81. Available from: https://pubmed.ncbi.nlm.nih.gov/23809438/

30. Nicolae CM, Aho ER, Choe KN, Constantin D, Hu HJ, Lee D, et al. A novel role for the mono-ADP-ribosyltransferase PARP14/ARTD8 in promoting homologous recombination and protecting against replication stress. Nucleic Acids Res [Internet]. 2015 Mar 31 [cited 2025 Feb 14];43(6):3143–53. Available from: https://pubmed.ncbi.nlm.nih.gov/25753673/

31. Sethy C, Kundu CN. 5-Fluorouracil (5-FU) resistance and the new strategy to enhance the sensitivity against cancer: Implication of DNA repair inhibition. Biomed Pharmacother [Internet]. 2021 May 1 [cited 2025 Feb 14];137. Available from: https://pubmed.ncbi.nlm.nih.gov/33485118/

32. Wyatt MD, Wilson DM. Participation of DNA repair in the response to 5- fluorouracil. Cell Mol Life Sci [Internet]. 2009 Mar [cited 2025 Feb 14];66(5):788–99. Available from: https://pubmed.ncbi.nlm.nih.gov/18979208/

33. Pommier Y. Drugging topoisomerases: lessons and challenges. ACS Chem Biol [Internet]. 2013 Jan 18 [cited 2025 Feb 14];8(1):82–95. Available from: https://pubmed.ncbi.nlm.nih.gov/23259582/

34. Harrision D, Gravells P, Thompson R, Bryant HE. Poly(ADP-Ribose) Glycohydrolase (PARG) vs. Poly(ADP-Ribose) Polymerase (PARP) - Function in Genome Maintenance and Relevance of Inhibitors for Anti- cancer Therapy. Front Mol Biosci [Internet]. 2020 Aug 28 [cited 2025 Feb 14];7. Available from: https://pubmed.ncbi.nlm.nih.gov/33005627/

35. Rack JGM, Palazzo L, Ahel I. (ADP-ribosyl)hydrolases: structure, function, and biology. Genes Dev [Internet]. 2020 Mar 1 [cited 2025 Feb 14];34(5–6):263–84. Available from: https://pubmed.ncbi.nlm.nih.gov/32029451/

36. Liu C, Vyas A, Kassab MA, Singh AK, Yu X. The role of poly ADP-ribosylation in the first wave of DNA damage response. Nucleic Acids Res [Internet]. 2017 Aug 21 [cited 2025 Feb 14];45(14):8129–41. Available from: https://pubmed.ncbi.nlm.nih.gov/28854736/

37. Tentori L, Leonetti C, Scarsella M, Muzi A, Vergati M, Forini O, et al. Poly(ADP-ribose) glycohydrolase inhibitor as chemosensitiser of malignant melanoma for temozolomide. Eur J Cancer [Internet]. 2005 Dec [cited 2025 Feb 14];41(18):2948–57. Available from: https://pubmed.ncbi.nlm.nih.gov/16288862/

38. Matanes E, López-Ozuna VM, Octeau D, Baloch T, Racovitan F, Dhillon AK, et al. Inhibition of Poly ADP-Ribose Glycohydrolase Sensitizes Ovarian Cancer Cells to Poly ADP-Ribose Polymerase Inhibitors and Platinum Agents. Front Oncol [Internet]. 2021 Oct 27 [cited 2025 Feb 14];11. Available from: https://pubmed.ncbi.nlm.nih.gov/34778062/

39. Li J, Koczor CA, Saville KM, Hayat F, Beiser A, McClellan S, et al. Overcoming Temozolomide Resistance in Glioblastoma via Enhanced NAD+ Bioavailability and Inhibition of Poly-ADP-Ribose Glycohydrolase. Cancers (Basel) [Internet]. 2022 Aug 1 [cited 2025 Feb 14];14(15). Available from: https://pubmed.ncbi.nlm.nih.gov/35892832/

40. Fabbrizi MR, Nickson CM, Hughes JR, Robinson EA, Vaidya K, Rubbi CP, et al. Targeting OGG1 and PARG radiosensitises head and neck cancer cells to high-LET protons through complex DNA damage persistence. Cell Death Dis [Internet]. 2024 Feb 1 [cited 2025 Feb 14];15(2). Available from: https://pubmed.ncbi.nlm.nih.gov/38368415/

41. Zhou X, Hao Q, Lu H. Mutant p53 in cancer therapy—the barrier or the path. J Mol Cell Biol [Internet]. 2019 Apr 1 [cited 2025 Feb 14];11(4):293–305. Available from: 10.1093/jmcb/mjy072

42. Sun Y, Chen J, Huang S yin N, Su YP, Wang W, Agama K, et al. PARylation prevents the proteasomal degradation of topoisomerase I DNA-protein crosslinks and induces their deubiquitylation. Nature Communications 2021 12:1 [Internet]. 2021 Aug 18 [cited 2025 Feb 14];12(1):1–16. Available from: https://www.nature.com/articles/s41467-021-25252-9

43. Finan JM, Niro R Di, Park SY, Jeong KJ, Hedberg MD, Smith A, et al. The polymeric fluoropyrimidine CF10 overcomes limitations of 5-FU in pancreatic ductal adenocarcinoma cells through increased replication stress. Cancer Biol Ther [Internet]. 2024 Dec 31 [cited 2025 Feb 14];25(1). Available from: https://www.tandfonline.com/doi/abs/10.1080/15384047.2024.2421584

44. Haber AO, Jain A, Mani C, Nevler A, Agostini LC, Golan T, et al. AraC- FdUMP[10] (CF10) is a next generation fluoropyrimidine with potent antitumor activity in PDAC and synergy with PARG inhibition. Mol Cancer Res [Internet]. 2021 Apr 1 [cited 2025 Feb 14];19(4):565. Available from: https://pmc.ncbi.nlm.nih.gov/articles/PMC9013283/

45. Wang JQ, Tang Y, Li QS, Xiao M, Li M, Sheng YT, et al. PARG regulates the proliferation and differentiation of DCs and T cells via PARP/NF-κB in tumour metastases of colon carcinoma. Oncol Rep [Internet]. 2019 May 1 [cited 2025 Feb 14];41(5):2657–66. Available from: https://pubmed.ncbi.nlm.nih.gov/30864743/

46. Marques M, Jangal M, Wang LC, Kazanets A, da Silva SD, Zhao T, et al. Oncogenic activity of poly (ADP-ribose) glycohydrolase. Oncogene [Internet]. 2019 Mar 21 [cited 2025 Feb 14];38(12):2177–91. Available from: https://pubmed.ncbi.nlm.nih.gov/30459355/

47. Yu M, Chen Z, Zhou Q, Zhang B, Huang J, Jin L, et al. PARG inhibition limits HCC progression and potentiates the efficacy of immune checkpoint therapy. J Hepatol [Internet]. 2022 Jul 1 [cited 2025 Feb 14];77(1):140–51. Available from: https://pubmed.ncbi.nlm.nih.gov/35157958/

48. Study Details | A Study of PARG Inhibitor ETX-19477 in Patients With Advanced Solid Malignancies | ClinicalTrials.gov [Internet]. [cited 2025 Feb 14]. Available from: https://clinicaltrials.gov/study/NCT06395519

49. Study Details | A Study of PARG Inhibitor IDE161 in Participants with Advanced Solid Tumors | ClinicalTrials.gov [Internet]. [cited 2025 Feb 14]. Available from: https://clinicaltrials.gov/study/NCT05787587

