## Supplementary Figures for "Loss-of-Function genetic Screen Unveils Synergistic Efficacy of PARG Inhibition with Combined 5-Fluorouracil and Irinotecan Treatment in Colorectal Cancer"

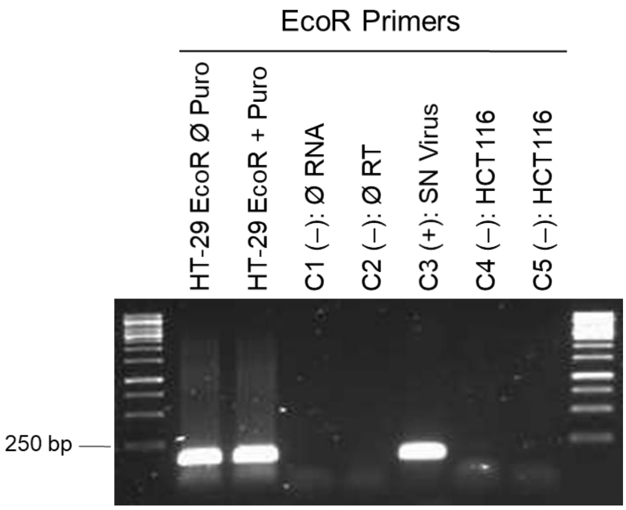


**Supplementary Figure S1. Validation of EcoR introduction into the HT29 cell line.** PCR amplification of the EcoR receptor (250 bp) was performed, and product was visualized in 1% agarose gel. The first two lanes correspond to EcoR infected HT29 cells before and after puromycin selection as indicated. Negative controls include the HCT116 cell line not infected with EcoR (C4-C5) and controls without RNA or retrotranscriptase (C1-C2, respectively). A viral supernatant containing infectious EcoR viral particles was used as a positive control (C3).

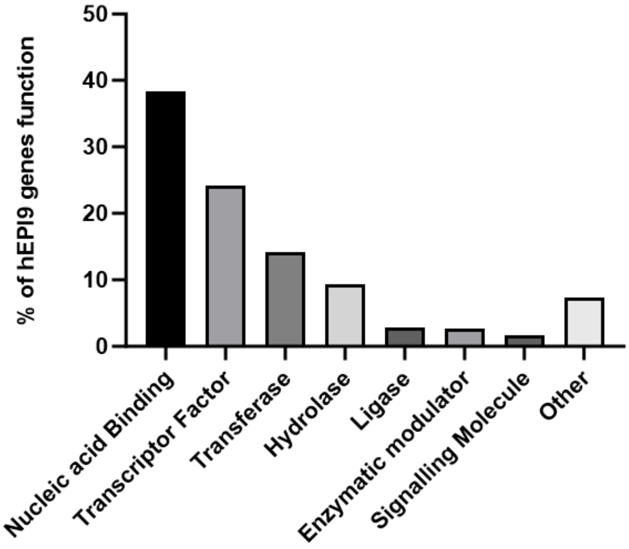


**Supplementary Figure S2. Biological function of hEPI9 shRNA library genes.** The bar plot shows the percentage of genes in each category.

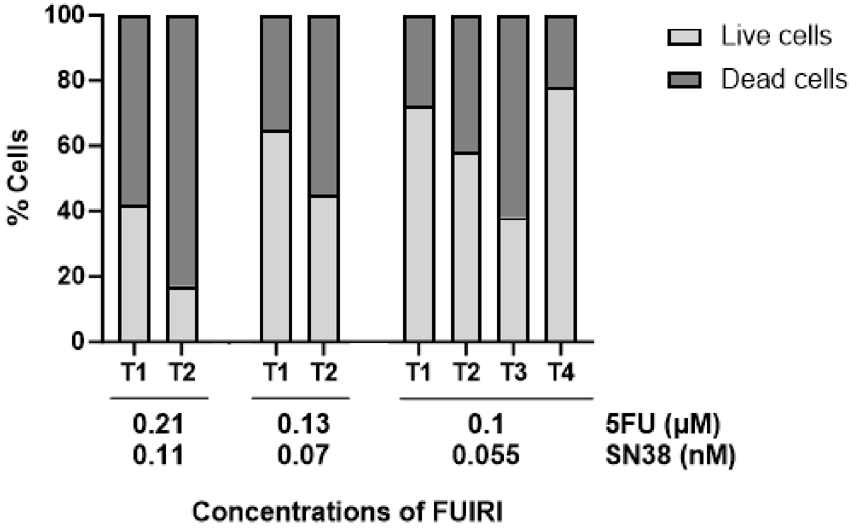


**Supplementary Figure S3. Determination of $IC_{20}$ dose of FUIRI treatment.** HT29 cells were treated with different doses of 5FU and SN38 combination maintained for 4 consecutive cycles. The percentage of live and dead cells was assessed using DiOC and DAPI staining and analyzed by flow cytometry. At high doses, only two treatment cycles were feasible due to increased cell mortality. The FUIRI $IC_{20}$ dose (0.1 uM 5FU + 0.055 nM SN38) resulted in 80% cell viability (light grey). The bars in the plot represent the mean of three independent experiments.

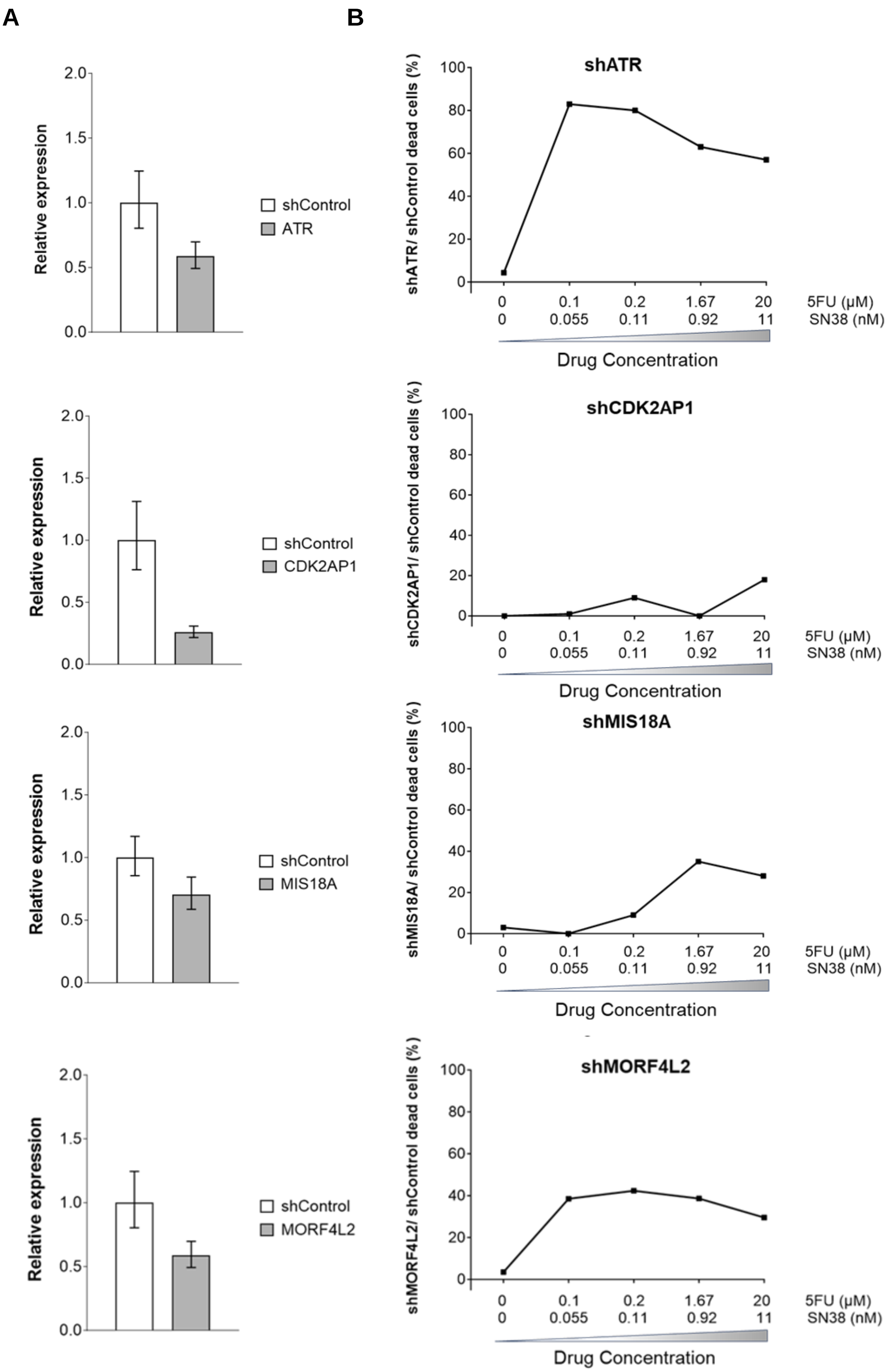
A
B
Relative expression
2.0
1.5
1.0
0.5
0.0
shControl
ATR
CDK2AP1
MIS18A
MORF4L2
shATR
shCDK2AP1
shMIS18A
shMORF4L2
shATR/ shControl dead cells (%)
shCDK2AP1/ shControl dead cells (%)
shMIS18A/ shControl dead cells (%)
shMORF4L2/ shControl dead cells (%)
100
80
60
40
20
0
0 0.1 0.2 1.67 20 5FU (µM)
0 0.055 0.11 0.92 11 SN38 (nM)
Drug Concentration

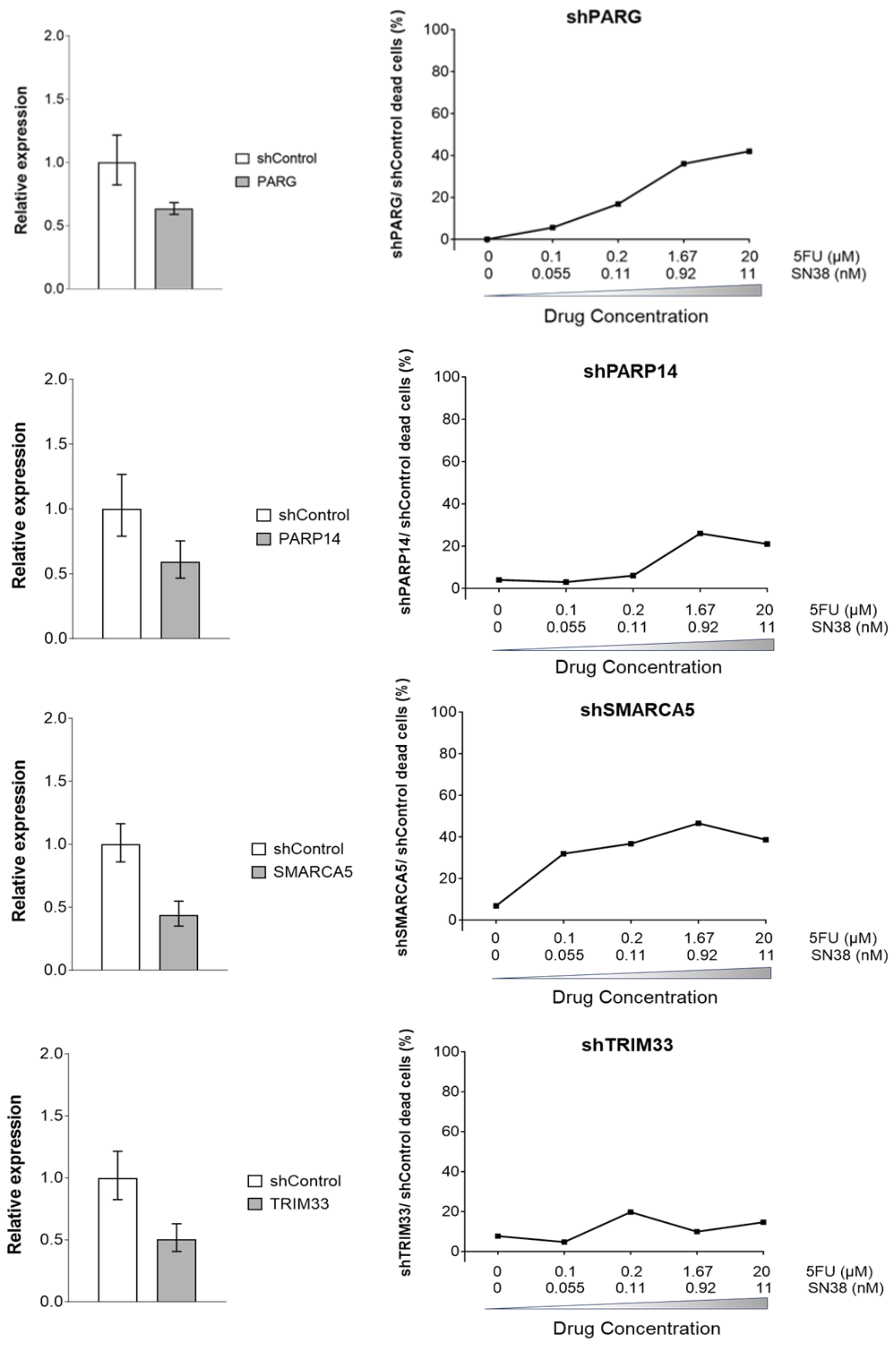


**Supplementary Figure S4. Individual Validation of LOF candidate genes that conferred sensitivity to FUIRI treatment.** A) HT-29 EcoR cells were infected with the most effective shRNA of each candidate gene and with the shRNA control. Candidate gene relative expression was measured by RT-qPCR, using the β-actin gene as a reference, and quantified using the $2^{(-\Delta\Delta Ct)}$ method. Bar graphs represent the mean expression levels of the specified genes relative to the shControl ± SD of at least 3 independent experiments. (B) FUIRI response based on the expression of selected genes. HT29 cells infected with the specified shRNAs or shControls were treated with different concentrations of FUIRI as indicated. Sensitivity to treatment was assessed by quantifying dead cells using DiOC6 and DAPI staining followed by flow cytometry. Graphs show the average of dead HT29 cells infected with the indicated shRNA relative to HT29 shControl cells of three independent experiments. The results are represented as percentages.

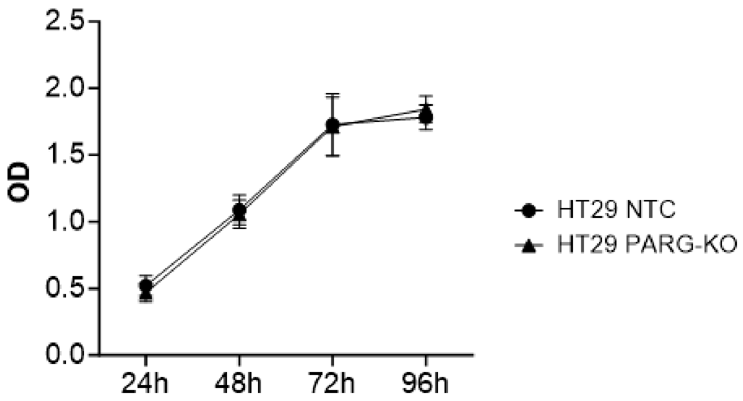


**Supplementary Figure S5. Proliferation of HT29 PARG-KO and NTC cells.** Cell proliferation was monitored over 96 hours to assess potential cell growth differences between NTC and PARG KO cells using propidium iodide staining. The results are presented as the mean optical density (OD) ± SD from three independent experiments.

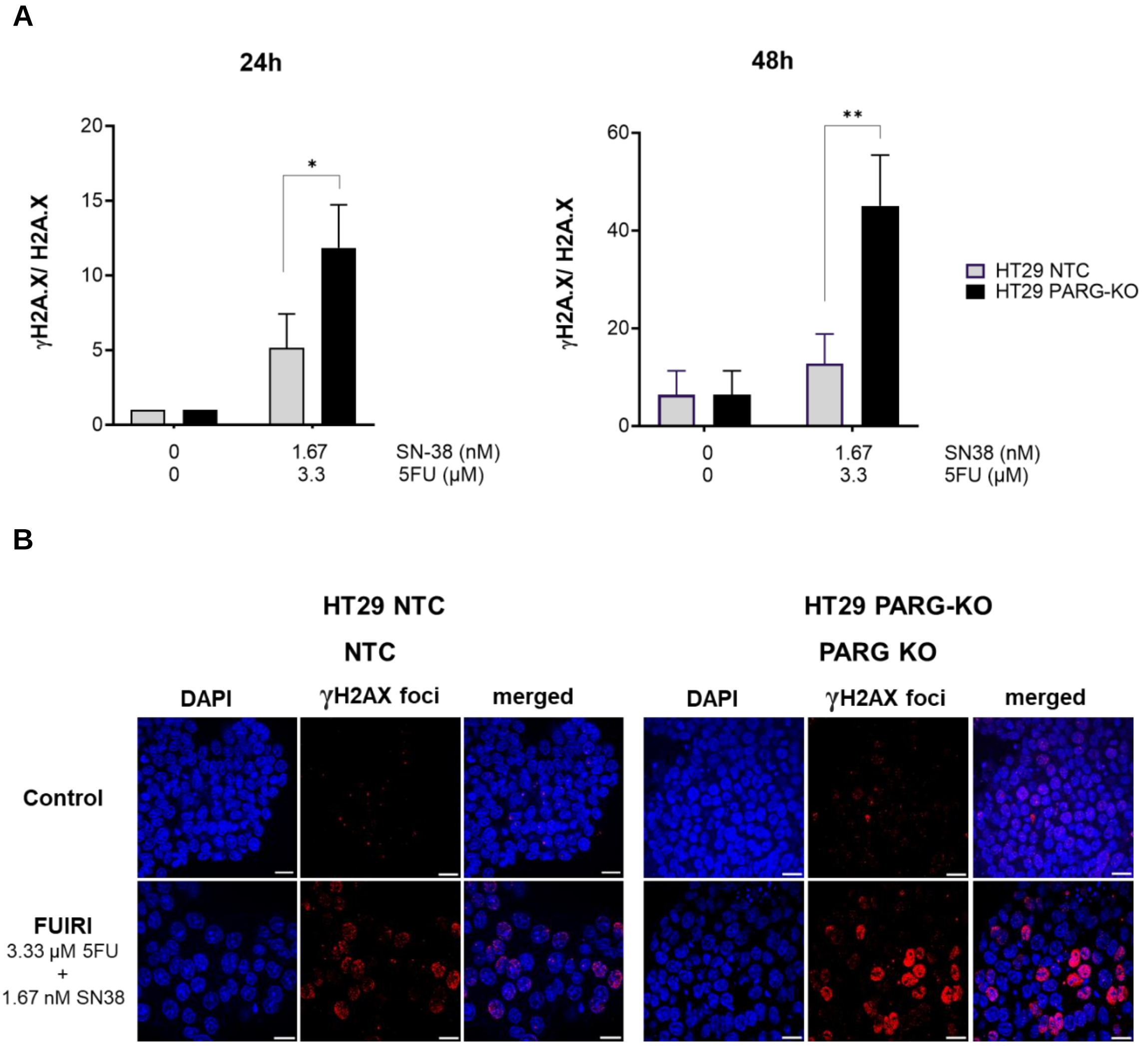


**Supplementary Figure S6**. **FUIRI induced DNA damage in HT29 NTC and PARG-KO cells.** A) HT29 NTC and PARG KO cells were treated with FUIRI for 24 hours. DNA damage was assessed immediately after treatment (left graph) or following 48 hours of recovery (right graph) by measuring γH2A.X protein levels via Western blot (WB), using the non-phosphorylated form of H2A.X as a normalizer. Bar graphs show the mean γH2A.X/H2A.X ratio ± SD from at least three independent experiments. B) Representative images of DNA damage, visualized by Immunofluorescence (IF) in cells treated with FUIRI. DNA damage was evaluated by detecting γH2AX foci in both HT29 NTC and PARG KO cells after 24 hours of FUIRI treatment and 48 hours of recovery. After fixation, nuclei were stained with DAPI (blue), and DNA damage (γH2AX foci) was detected with an anti-γH2A.X antibody (red) for IF analysis. Objective lens: 63X immersion oil. Scale bar: 30 μm. Differences in DNA damage were statistically analyzed using Student's t-test (*$p < 0.05$; **$p < 0.01$).

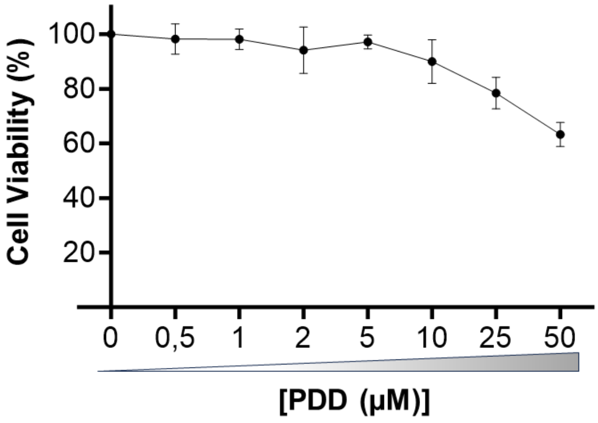


**Supplementary Figure S7. PDD activity as a single agent.** HT29 cells were treated for 72 hours with different concentrations of the PARG inhibitor PDD, as indicated. Cell proliferation was assessed using the MTT assay. The graph represents the mean percentage of viable treated cells relative to non-treated controls ± SD.

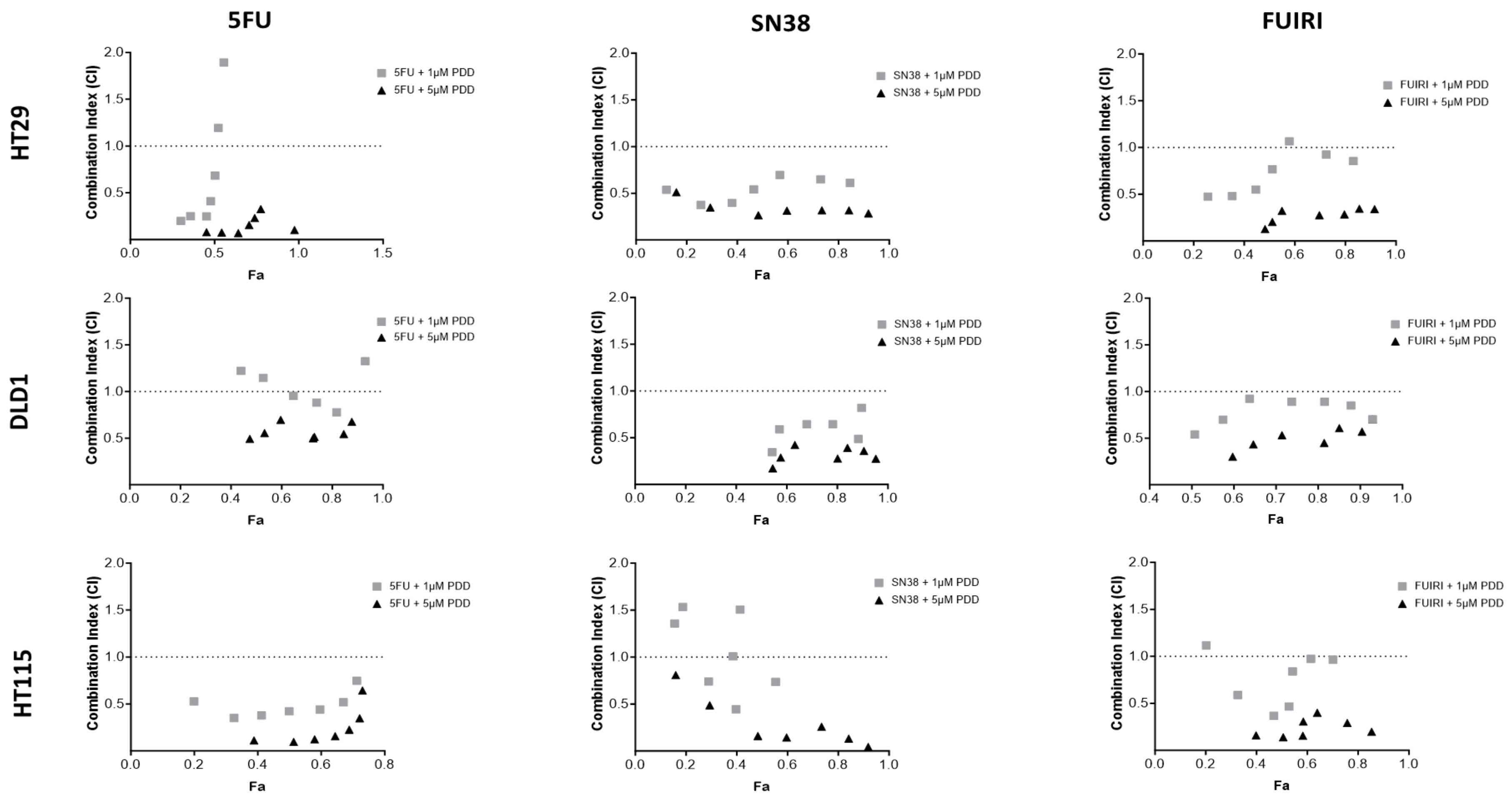


**Supplementary Figure S8. Combination index (CI) of 5FU, SN38, and FUIRI with the PARG inhibitor PDD.** HT29, DLD1, and HT115 cell lines were treated for 72 hours with different concentrations of 5FU, SN38, and FUIRI combination with either 1 µM and 5 µM of PDD. The graphs display the CI-Fa (fraction affected) plot, based on the method proposed by Chou and Talalay. The combination index (CI) was calculated using CompuSyn software, which quantitatively defines drug interactions as synergistic (CI < 1), additive (CI = 1), or antagonistic (CI > 1).

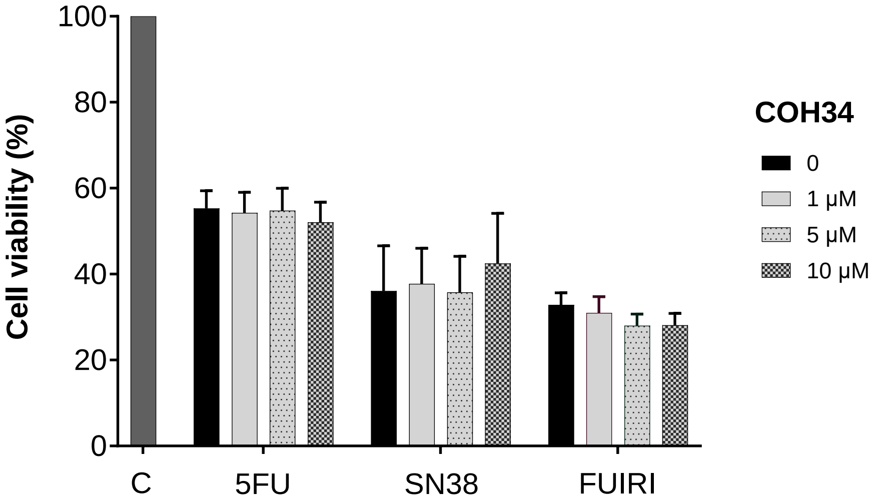


**Supplementary Figure S9. Effect of COH34 PARG inhibitor in combination with 5FU, SN38 and FUIRI in HT29 cell line.** HT29 cells were treated for 72 hours with 5FU (12 μM), SN38 (30 nM), or FUIRI (12 μM 5FU + 30 nM SN38) in combination with 0 μM, 1 μM, 5 μM, and 10 μM of COH34. Cell viability was assessed using the MTT assay. Bar graphs represent the mean percentage of viable treated cells relative to non-treated controls ± SEM. Statistical differences between treatments were analyzed using the Student's t-test. C = control.

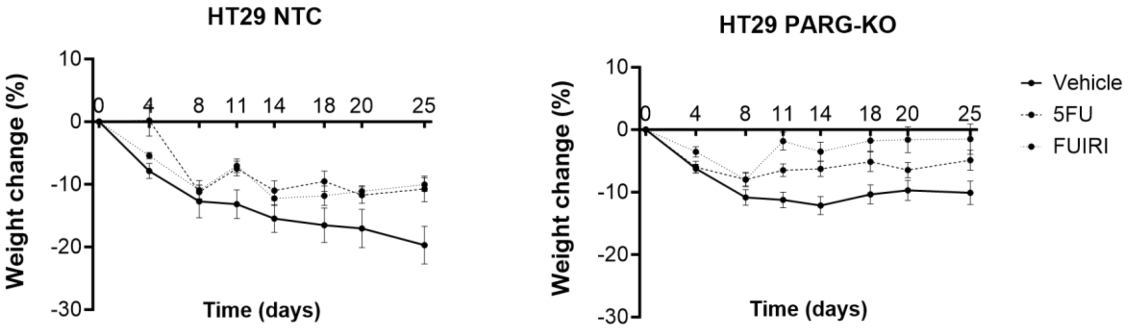


**Supplementary Figure S10**. ***In vivo* toxicity of 5FU and FUIRI.** Balb/c mice bearing HT29 NTC and PARG-KO tumors were treated once a week for 25 days with vehicle (PBS), 5FU (50 mg/kg), or FUIRI (50 mg/kg 5FU + 50 mg/kg CPT-11). Toxicity was evaluated by monitoring body weight loss, calculated as the percentage change relative to day 0. Weight measurements were taken twice a week. Graphs represent the mean percentage of weight change for ten mice per group at each time point ± SEM. A weight loss exceeding 20% was considered indicative of treatment toxicity.

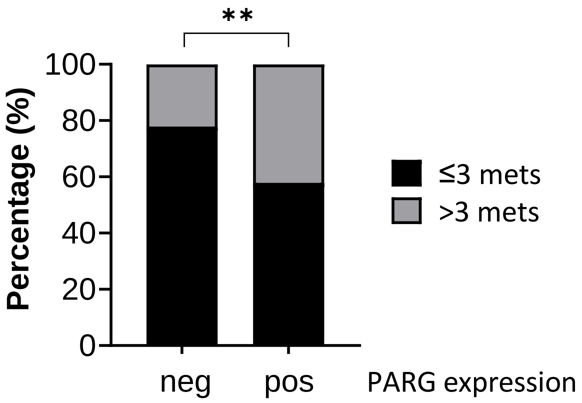


**Supplementary Figure S11. PARG expression and liver metastases.** Bar graphs showing the proportion of patients with fewer than three or at least three liver metastases categorized according to PARG expression (negative or positive) by IHC staining. Statistical differences were analyzed using Fisher's exact test (**p = 0.01).
