## Supplementary Tables for "Loss-of-Function genetic Screen Unveils Synergistic Efficacy of PARG Inhibition with Combined 5-Fluorouracil and Irinotecan Treatment in Colorectal Cancer"

**Supplementary Table ST1. Molecular characteristics of CRC cell lines.**

**A**

| Gene Name | DLD1 | HT115 | HT29 |
|---|---|---|---|
| ***ATM*** | | p.D130Y | |
| ***ATR*** | p.I1851V | p.R1015q | |
| ***ATRX*** | p.L629R | p.R1138I<br>P.K869N | |
| ***BRAF*** | | p.R354Q | p.V600E |
| ***BRCA1*** | | p.S763F<br>p.K1732R<br>p.E577* | |
| ***BRCA2*** | | p.E2258K<br>p.S2052*<br>p.L29I | |
| ***CDH1*** | | p.E648* | |
| ***CDH11*** | p.L624I<br>p.G242S<br>p.G96* | | |
| ***CDK12*** | | p.D214Y | |
| ***CHEK2*** | p.A290D | p.N333T | |
| ***ERCC1*** | p.P33H | | |
| ***ERCC4*** | c.2747del | p.D231G | |
| ***FANCA*** | p.G110C<br>p.E1178* | | |
| ***FANCD2*** | p.R1273Q | | |
| ***KRAS*** | p.G13D | | |
| ***MLH1*** | p.A120S | | |
| ***MSH6*** | p.Y469F | p.E1322*<br>p.R911Q | |
| ***MTOR*** | | p.A695V | |
| ***MUTYH*** | p.Q138R | | |
| ***NBN*** | p.A710S | | |
| ***PALB2*** | p.G964V | p.E554K | |
| ***PIK3CA*** | p.E545K<br>p.D549N | p.R88Q<br>p.E321D<br>p.R770Q | p.P449T |
| ***RAD21*** | | p.R450H | |
| ***SMARCB1*** | p.T390A | | |
| ***TP53*** | p.S241F | p.R213* | p.R273H* |

**B**

| Cell Line | MSI status | CIMP | CMS |
|---|---|---|---|
| **DLD1** | MSI | CIMP + | CMS1 |
| **HT115** | MSS | CIMP - | CMS3 |
| **HT29** | MSS | CIMP + | CMS3 |

A) Mutational profile of DLD1, HT115 and HT29 cell lines. Mutations in the most important DNA repair genes and key CRC-associated genes were verified using Cosmic and Cellosaurus databases. Only missense and nonsense mutations classified according to TIER consensus are presented. B) Additional molecular cell line characteristics. Microsatellite Status (MSI = Microsatellite instable; MSS = Microsatellite stable), CpG island methylator phenotype (CIMP + = positive; CIMP - = negative) and Consensus Molecular Subtype (CMS).

**Supplementary Table ST2. Primers used in this study.**

| Name | Strand | Sequence (5' to 3') |
|---|---|---|
| **EcoR receptor** | F<br>R | GGGTTTATGCCCTTTGGATT<br>CACGCCAAAGTACGCTATGA |
| **ATR** | F<br>R | GCCCAGACAAGCATGATCCAG<br>AAGATGATGACCACACTGAGA |
| **CDK2AP1** | F<br>R | CAGCTGCTCAGTGACTACG<br>TCCCCAGCTCTTCAATGATG |
| **MIS18A** | F<br>R | GCTGGTGTTCCTGTGCT<br>GATAGCTTCTGTTCCTTATCCACA |
| **MORF4L2** | F<br>R | GTGCGTATTTGCCTGAAGAAG<br>TCCTCACTATACAGCAAACTTAGC |
| **PARG** | F<br>R | TGCTGAGACATATCGTTGGTC<br>GAGGTAGCGTCTGAAGTGAA |
| **PARP14** | F<br>R | ACTTGAACACATACACTGCCA<br>TTCTGCTGCTTCATATCACTCC |
| **SMARCA5** | F<br>R | GTGGTCTTGGCATCAATCTTG<br>AAAGCGGAACACTCTGACTG |
| **TRIM33** | F<br>R | TCCCAACACTACCAAATCCC<br>CTGCCTGAGCTCTTCTGAATC |
| **β-Actin** | F<br>R | TGAGCGCGGCTACAGCTT<br>TCCTTAATGTCACGCACGATTT |
| **PARG CRISPR** | F<br>R | ATTTCTTGCCTGGGAGCTGG<br>ACACAGGGCCCATAAACAGG |

All primers shown were used for gene expression analysis by RT-qPCR, except for the PARG CRISPR as these primers were used to confirm CRISPR/Cas9 gene editing by PCR amplification and by subsequent Sanger sequencing using the forward primer.

**Supplementary Table ST3. CRISPR/Cas9 guide sequences for Non-Targeting Control and PARG gene**

| Gene | gRNA Number Genescript | Sequence (5' to 3') | Gene Location |
|---|---|---|---|
| Control | NonTargeting Control Guide for Human | ACGGAGGCTAAGCGTCGCAA | - |
| PARG | crRNA1<br>crRNA2<br>crRNA3 | GTTCTTACCTCATCTTCCAC<br>TGCTATTCTGAAATACAATG<br>AAAGAATGGTGAGCGAACTG | Exon 5<br>Exon 7<br>Exon 6 |

CRISPR guides used to target PARG gene as well as NTC are represented in this table

**Supplementary Table ST4. Top 24 depleted genes obtained in the LOF screening**

| Gene Name | logFC-sh1 | logFC-sh2 | logFC-sh3 | logFC-sh4 | logFC-sh5 | logFC-sh6 | logFC-sh7 | ogFC-sh8 | mean (<0) | P Value Mixed | FDR Mixed |
|---|---|---|---|---|---|---|---|---|---|---|---|
| **ATR** | 0,037 | -1,925 | -1,064 | -0,637 | -1,060 | -0,538 | -0,668 | -0,767 | -0,95 | 0,001 | 0,456 |
| **SMARCA5** | -0,568 | -0,574 | -0,893 | 0,050 | -0,481 | -1,645 | -0,647 | 1,095 | -0,80 | 0,027 | 0,464 |
| **ATM** | -0,708 | 0,009 | -0,288 | -0,591 | -1,146 | 0,028 | -1,077 | -0,376 | -0,70 | 0,038 | 0,496 |
| **ATXN7L3** | -0,752 | -0,960 | -0,745 | -0,365 | -0,065 | 0,143 | 0,272 | -1,222 | -0,68 | 0,004 | 0,464 |
| **TOX4** | -0,911 | -0,878 | -0,911 | 0,282 | -0,240 | -0,208 | -0,349 | 0,216 | -0,58 | 0,031 | 0,480 |
| **BRD4** | -0,281 | -1,092 | -0,451 | -0,613 | -0,521 | -0,842 | -0,166 | 0,279 | -0,57 | 0,026 | 0,464 |
| **ING3** | -0,411 | 0,207 | -0,354 | 0,420 | -0,557 | -0,731 | -1,102 | -0,165 | -0,55 | 0,071 | 0,516 |
| **MIS18A** | -0,005 | 0,030 | -0,959 | -0,513 | -1,152 | -0,506 | -0,362 | -0,267 | -0,54 | 0,082 | 0,516 |
| **SCMH1** | 0,685 | -0,798 | -0,858 | -0,301 | -0,929 | -0,160 | 0,365 | -0,146 | -0,53 | 0,026 | 0,464 |
| **TBL1XR1** | -1,240 | -0,421 | -0,640 | -0,489 | -0,373 | -0,353 | 0,324 | -0,044 | -0,51 | 0,071 | 0,516 |
| **TADA3** | -0,749 | 0,345 | -0,494 | -0,222 | -0,117 | -0,565 | -0,891 | 1,425 | -0,51 | 0,021 | 0,464 |
| **PARG** | -0,072 | -0,629 | -0,635 | -0,219 | -0,450 | -0,471 | 0,082 | -0,977 | -0,49 | 0,07 | 0,516 |
| **FBXL19** | -0,561 | -0,799 | 0,514 | -0,175 | 0,401 | -0,186 | -0,586 | -0,399 | -0,45 | 0,084 | 0,516 |
| **KDM5A** | -0,237 | 0,235 | -0,389 | -1,321 | -0,227 | -0,391 | -0,015 | 0,435 | -0,43 | 0,02 | 0,464 |
| **PRKAA1** | -0,663 | -0,108 | -0,720 | -0,842 | -0,291 | -0,038 | -0,308 | 0,176 | -0,42 | 0,056 | 0,516 |
| **BRCA1** | 0,056 | -0,301 | -0,514 | 0,445 | -0,560 | -0,019 | -0,801 | -0,308 | -0,42 | 0,009 | 0,464 |
| **GLYR1** | -0,179 | -0,289 | 0,419 | -0,409 | -0,254 | -0,253 | -0,542 | -0,970 | -0,41 | 0,046 | 0,516 |
| **PARP14** | -0,428 | -0,231 | -0,772 | 0,658 | -0,464 | -0,074 | 0,001 | -0,270 | -0,37 | 0,023 | 0,464 |
| **CDK2AP1** | 2,001 | -0,761 | -0,737 | 0,105 | -0,282 | -0,199 | -0,093 | -0,123 | -0,37 | 0,029 | 0,464 |
| **TRIM33** | -0,282 | 0,575 | -0,182 | -0,236 | -0,577 | 0,224 | -0,533 | -0,357 | -0,36 | 0,054 | 0,516 |
| **MECOM** | 0,232 | -0,223 | -0,246 | -0,683 | -0,310 | 0,255 | -0,476 | -0,161 | -0,34 | 0,007 | 0,464 |
| **INTS12** | -0,259 | -0,631 | -0,090 | -0,272 | -0,058 | 0,392 | -0,525 | 0,685 | -0,31 | 0,066 | 0,516 |
| **AFF4** | -0,254 | -0,037 | -0,441 | 0,576 | -0,397 | -0,080 | -0,136 | -0,593 | -0,28 | 0,034 | 0,483 |
| **MORF4L2** | 0,056 | -0,145 | 0,083 | -0,100 | -0,088 | -0,587 | -0,314 | -0,129 | -0,23 | 0,061 | 0,516 |

24 top drop-out genes identified after four consecutive FUIRI treatments showing the logFC values for each shRNA individually (sh1 to sh8), the mean of logFC of the 8 shRNA, p-values < 0.1 combined from 8 shRNA per gene and FDR values < 0.5.
